# MERTK inhibition cooperates with immunomodulatory cyclophosphamide to induce CXCL9⁺ monocyte-macrophage programming and durable anti-tumor immunity in triple negative breast cancer

**DOI:** 10.64898/2026.02.04.703648

**Authors:** Alex J. Smith, Zachary Schrank, Nan Guan, Diego A. Pedroza, Sebastian Calderon, Xueying Yuan, Na Zhao, Zoe Gabriel, Yang Gao, Charlotte Rivas, Fengshuo Liu, Jonathan S. Serody, Charles M. Perou, H. Shelton Earp, Jeffrey M. Rosen

## Abstract

Triple-negative breast cancer (TNBC) has high rates of recurrence despite chemotherapy and immune checkpoint blockade (ICB). Tumor-associated macrophages (TAMs) can either suppress or support anti-tumor immunity, but the mechanisms governing these states and therapeutic targets remain unclear. Here, integrating public scRNAseq datasets with TNBC cohorts, we identify a prognostic myeloid signature defined by CXCL9hi/C1Qlow TAM programs, associated with improved survival and increased lymphocyte activation pathways. Using immunocompetent p53-null syngeneic TNBC models spanning basal-like (2153L) and claudin-low (T12) subtypes, we show that immunomodulatory cyclophosphamide (CTX) reprograms hematopoiesis toward the monocytic lineage and induces an interferon (IFN) conditioned tumor milieu that supports CXCL9⁺ monocyte-derived macrophages (Mo.Macs) in basal-like disease. Combining CTX with the next generation MERTK-selective inhibitor UNC2371 (MRX-2843) drives complete remissions in both models, but durable long-term responses occurred selectively in the basal-like subtype model. The expansion of antigen-presenting CXCL9⁺ Mo.Macs and reduction of C1q⁺ phagocytic TAMs are observed in responding tumors. Mechanistically, MERTK inhibition relieves MAPK/SOCS1 mediated restraint of IFN signaling driving a positive feedback loop of IRF7/STAT1/IRF1 driven CXCL9 induction. Functionally, tumor control requires CXCL9-CXCR3 dependent CD4⁺ T cell recruitment, accumulation of stem-like memory CD4⁺ T cells, and germinal center like immune organization in tumor-draining lymph nodes. PD-1 blockade further increases durability, preventing recurrence in most treated basal-like tumors. Together, these findings define an IFN licensed, MERTK regulated myeloid checkpoint that can be therapeutically targeted to convert suppressive TNBC microenvironments into durable adaptive immunity, supporting clinical translation of CTX + MRX-2843 based combinations in basal-like TNBC.

**Significance:** Suppressive myeloid programing limits effective adaptive immune engagement in TNBC usually resulting in ICB treatment resistance and tumor recurrence. This study identifies a therapeutically actionable myeloid interferon checkpoint in which MERTK inhibition stabilizes CXCL9⁺ monocyte-macrophage programming to promote CD4⁺ T cell dependent immune memory and durable tumor control in basal-like TNBC.

## INTRODUCTION

Breast cancer is the second leading cause of cancer-related deaths in women, and triple negative breast cancer (TNBC) has the worst prognosis (1, 2). TNBC disproportionately affects African American and young women, with a five-year survival rate of 77% (3, 4). This high mortality reflects tumor heterogeneity (5), and metastatic propensity (6). TNBC is now treated with immune checkpoint blockade (ICB) combined with multi-agent chemotherapy in the early-stage neoadjuvant setting (7, 8). However, the treatment landscape for TNBC may evolve soon as antibody drug conjugates continue to progress through clinical trials (9, 10). While some patients achieve pathological complete response (pCR), many retain residual disease leading to recurrence. The tumor microenvironment (TME) can be both predictive and prognostic for ICB response (11), but high cellular variability and substantial therapeutic resistance necessitate further study.

Tumor-infiltrating lymphocytes (TILs) predict response to chemotherapy and ICB in TNBC (12–14). In contrast, suppressive myeloid infiltration, including tumor associated macrophages (TAMs), correlates with poor prognosis (15). Pro-tumoral TAMs promote chemoresistance (16), angiogenesis (17), and metastasis (18–20), and coordinate lymphocyte-devoid ‘immune cold’ TMEs (21–24). Conversely, TAMs can also recruit and activate lymphocytes through cytokine release (25) and immune co-stimulatory protein expression (26), leading to enhanced anti-tumor immune responses.

Previously, our laboratory has extensively characterized a collection of preclinical p53null Balb/c murine tumors. This tumor bank was created by mammary fat pad transplantation from p53null donors into WT recipients (27). As p53 is mutated in over 90% of TNBC (28), genetic and TME characterization revealed that the tumor bank recapitulated clinical TNBC subtypes (29, 30) including basal-like and claudin-low (CL). This collection of tumors is a unique tool to model TNBC cancer cells (31, 32), tumor infiltrating myeloid cell heterogeneity (33), identify novel therapeutic targets (34–36), and mechanisms/treatment of metastasis (37–39).

With these models we have shown that immunomodulatory cyclophosphamide (CTX) combined with CSF1R inhibition and aPD1 drives adaptive memory and long-term durable responses (LTRs) in CL models (40). These studies have led to the New START (NCT06959537) phase Ib clinical trial. However, CL tumors represent only 10-15% of TNBC, motivating the search for TAM-targeting agents effective in basal-like TNBC, which comprises ∼60-80% of cases (41).

The TAM receptor tyrosine kinase family (TYRO3, AXL, MERTK) signaling exhibits both physiologic and pathophysiologic consequences (42, 43). The most well-defined receptor function is efferocytosis, the receptor directed phagocytosis of apoptotic cells, drives tissue level homeostasis in an immune tolerant manner to prevent autoimmunity (44, 45). However in the harsh TME of cancers, that physiologic TAM RTK function is subverted to the advantage of the tumor decreasing adaptive immune function and enhancing a wound-healing milieu by supporting angiogenesis (46, 47) and decreasing interferon signaling (48). Conversely, the TAM RTKs are often induced and overexpressed in tumor cells where they provide a therapy resistance survival signal by regulating ferroptosis (49), or in the case of AXL, enhance proliferation or epithelial to mesenchymal transition (50, 51). Due to their pro-tumor roles, TAM receptors are actively targeted in various cancer types clinically (52–54).

Here we investigated a next-generation TAM receptor inhibitor, UNC2371, which is now entering clinical evaluation as MRX-2843. Compared to previous iterations of the compound, MRX-2843 showed increased bioavailability as well as MERTK specificity. MRX-2843 has been shown to increase overall survival in preclinical models of acute myeloid leukemia (55), acute lymphoblastic leukemia (56), EGFR inhibitor-resistant non-small cell lung cancer (NSCLC) (57), glioblastoma (58), and Ewing sarcoma (59). Multiple clinical trials have been developed to investigate MRX-2843 in combination with Osimertinib for NSCLC (NCT04762199), refractory metastatic solid tumors (NCT03510104), and various subtypes of leukemia (NCT04946890, NCT04872478).

MRX-2843 has not been evaluated in models of TNBC and has yet to be coupled with immunomodulatory chemotherapies such as CTX. Here we show that when MRX-2843 is coupled with low-dose CTX it results in complete responses in both the 2153L basal-like and T12 CL models. We then identify TME differences between basal-like and CL tumors that are predictive of both preclinical and clinical response.

## RESULTS

### CXCL9⁺ and C1q⁺ TAM populations are prognostic in TNBC

Macrophage populations can be correlated with response in cancer patients. This is commonly attributed to the functional inability to recruit and activate lymphocytes. CXCL9^+^ has been shown to be an important chemokine secreted by myeloid cells to recruit and activate anti-tumor effector T cells (60–62), whereas C1q^+^ macrophages have been shown to correlate with a poor prognosis in multiple cancer types (63–65). To determine if these populations exist in human TNBC, if they are found in different proportions within different subtypes of TNBC, and if they correlate with T cell infiltration and response, we analyzed publicly available scRNAseq datasets. We could then compare these populations with our murine models, allowing for mechanistic studies of these populations.

Subclustering of the myeloid compartment in a publicly available single-cell atlas of ten human triple-negative breast cancers (GSE176078, (66–68) revealed CXCL9+ and C1q^+^ TAM populations (Figure 1A–B, S1A). CXCL9^+^ TAMs were associated with upregulated T/B cell activation pathways, while C1q^+^ TAMs correlated with upregulated organelle maintenance and T cell inhibitory pathways (Figure 1C, S1C). We then generated a 50-gene combined signature for these CXCL9^+^ and C1q^+^ TAM populations (Supplemental tables 1-2). Using The Cancer Genome Atlas (TCGA) breast cancer cohort, we found that TNBC patients with a CXCL9^+^ signature score above the median and C1q^+^ signature score below the median (CXCL9-high/C1q-low) had a greater 5-year overall survival than patients with CXCL9-low/C1q-high signature scores (Figure 1D). Both CXCL9^+^ and C1q^+^ TAMs were also present across p53null TNBC preclinical models. In p53null T12 (CL) and 2153L (basal-like) tumors, scRNAseq revealed that both models harbor CXCL9^+^ and C1q^+^ expressing TAMs, with T12 enriched for C1q^+^ TAMs and 2153L enriched for TILs and IFN signaling (Figure 1E-F, S1D-E). Comparison of PMNC populations between human and murine tumors revealed 2153L tumors closely model human TNBC (Figure 1G). Importantly, both type I and type II interferon (IFN) protein levels are higher at baseline in 2153L tumors, when compared to T12 tumors (Figure 1H).

**Figure 1.**
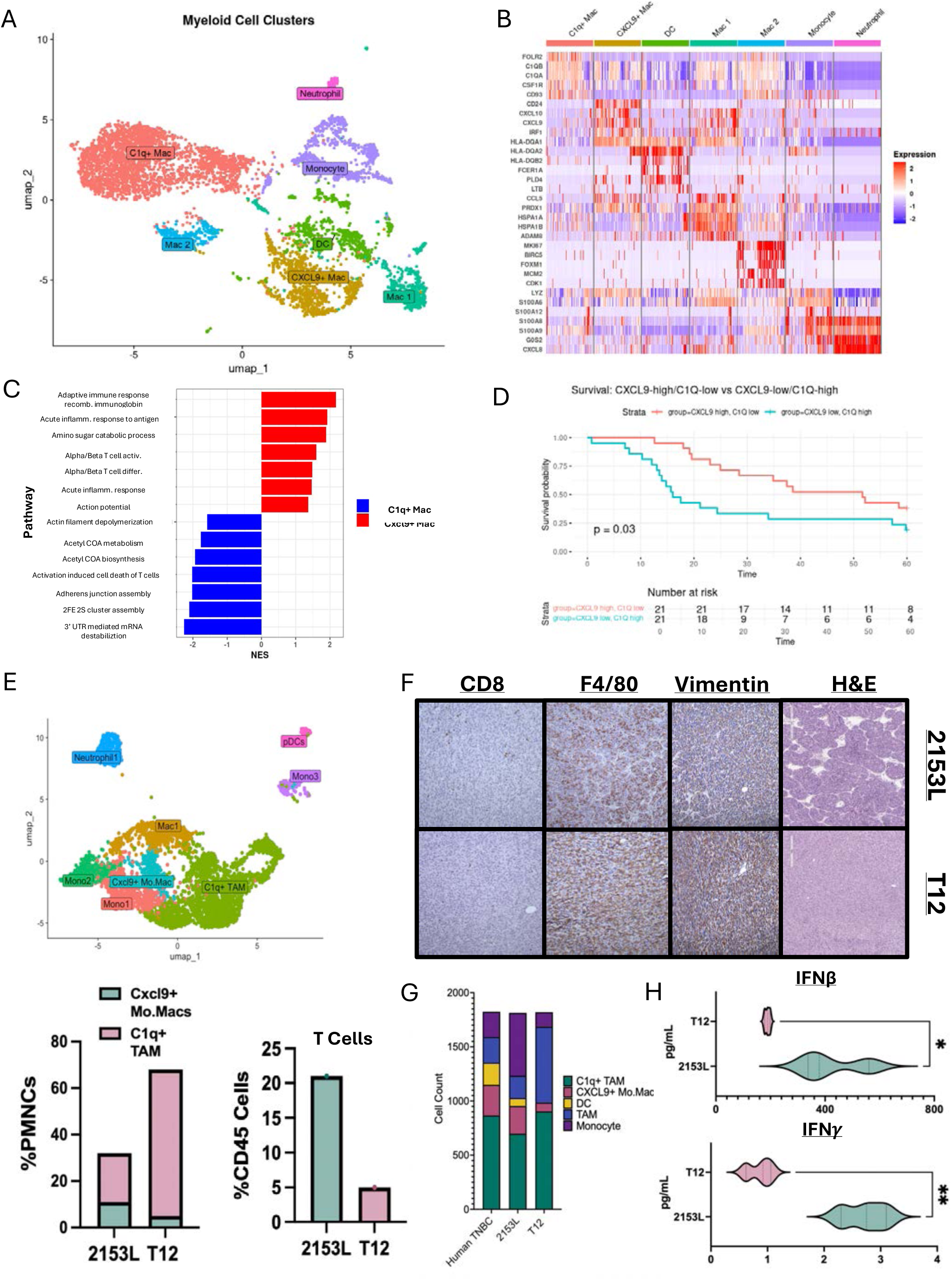
CXCL9+ and C1q+ macrophage populations are prognostic and predict a T Cell enriched or devoid tumor microenvironment. (A) Umap subclustering of PMNC populations from breast scRNAseq atlas. (B) Heatmap showing top DEG genes upregulated in each cluster C1qa, b, and c are upregulated in C1q+ population. Cxcl9 is upregulated in Cxcl9+ population. (C) GSEA of top 25 genes in Cxcl9+ population (Red) and C1q+ (Blue). (D) Survival KM plot comparing Cxcl9 high, C1q low signature (Red) to Cxcl9 low, C1q high signature (Blue). (E) Umap subclustering of untreated 2153L and T12 PMNC populations. (F) Cell population comparisons between mouse and human PMNC from scRNAseq datasets. (G) Protein quantification and comparison of IFNγ and IFNβ at baseline in 2153L and T12 tumors. (H) Comparison of Cxcl9^+^, C1q^+^, and T Cell populations between two murine models. (I) H&E and IHC staining for Vimentin, F4/80, and CD8 on 2153L and T12 p53null tumor models.

### CTX reprograms myelopoiesis towards monocytes and expands IFN-driven Mo.Macs in basal-like TNBC

The standard-of-care treatment for TNBC includes multiple neoadjuvant chemotherapies, which includes CTX (600mg/m^2^) with pembrolizumab, followed by surgical resection (69). Studies in mice have shown that 100 mg/kg of CTX is immunomodulatory due to clearance of suppressive immune cell populations via lymphodepletion (70–73). Furthermore, CTX treatment increases TILs and immune activation through IFN production due to its effect as a DNA alkylating agent (74–76). In this study, we compared the effects of immunomodulatory CTX on the TME and peripheral myelopoiesis within mice bearing 2153L or T12 tumors.

scRNAseq of CTX-treated tumors from both models at day 7 was subclustered to analyze phagocytic mononuclear cells (PMNC) and showed expansion of monocytes, TAMs, and monocyte-derived dendritic cells (Mo-DCs); the monocyte and TAM populations were validated independently with flow cytometry (Figure 2A, S1F). CTX increased C1q^+^ phagocytic TAMs in T12, while 2153L increased Ly6Chi monocytes and supported the differentiation of CXCL9^+^ Mo.Macs. This population expressed both antigen presentation genes and lymphocyte-activating gene pathways (Figure 2B–D, S1G-H). CTX induced DNA damage in 2153L tumors (Figure S2A). Interestingly, we observed MERTK to be strongly co-expressed with C1q (Figure S2B).

**Figure 2.**
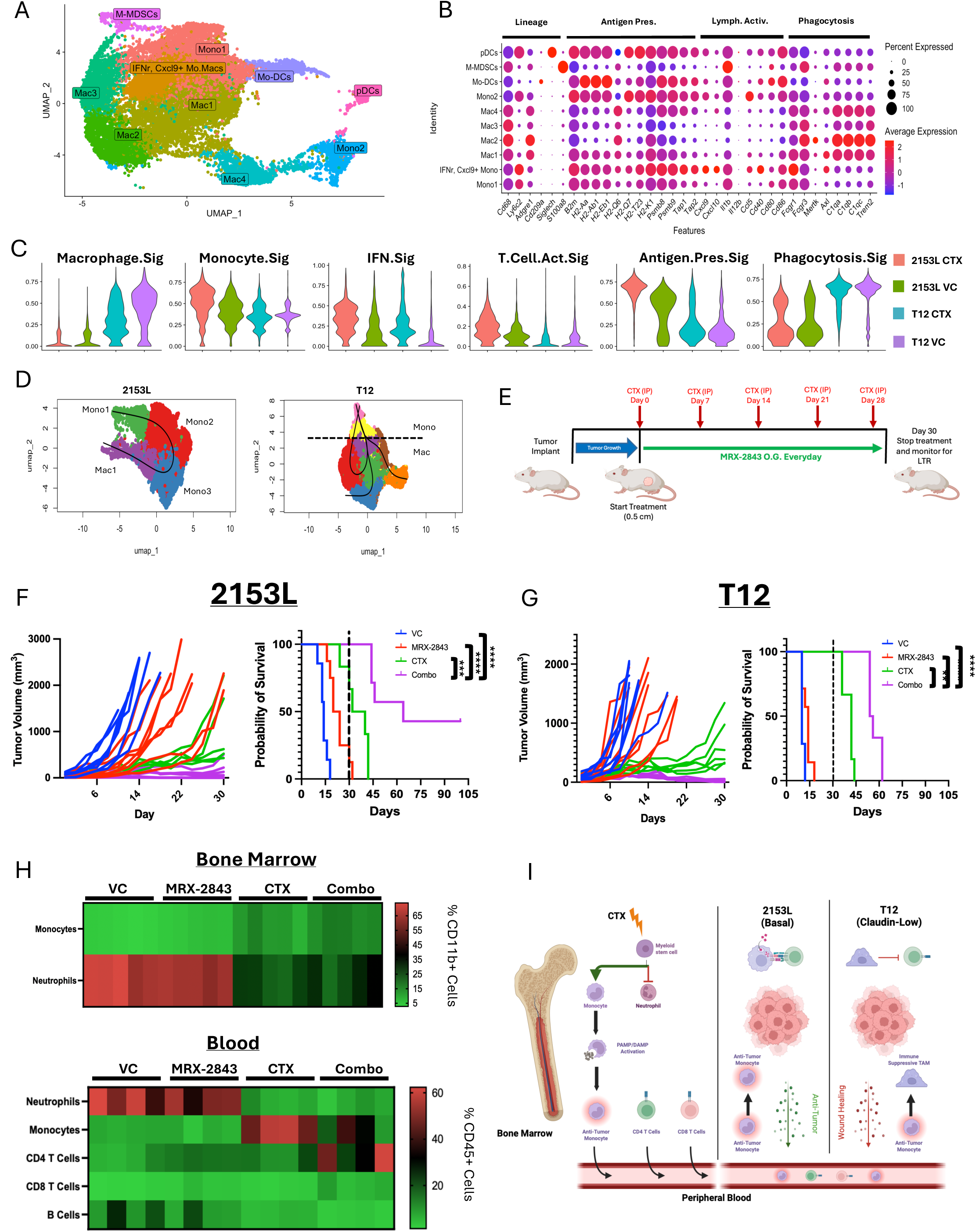
CTX reprograms myelopoiesis and increases IFN monocytes leading to long term durable responses in 2153L model. (A) Umap subclustering of PMNC from 2153L CTX treated, 2153L untreated, T12 CTX treated, and T12 untreated (Left to Right) scRNAseq datasets. (B) Comparison of PMNC population expression of lineage, antigen presentation, lymphocyte activation, and phagocytic gene markers within all 4 datasets. (C) UCell signature expression scoring of macrophage, monocyte, IFN, T cell activating, antigen presentation, and phagocytic markers for each dataset. (D) Slingshot pseudotime comparison of different monocyte differentiation in 2153L and T12. (E) Treatment strategy for CTX and MRX-2843 long-term response study. (F) 2153L tumor volume comparisons across treatments during 30 days of treatment. KM plot of overall survival for 2153L model. HR compared to VC n=7, MRX-2843 n=8 (HR = 0.079, p = 0.0026), CTX n=6 (HR = 0.019, p < 0.001), Combo n=7 (HR = 0.0014, p < 0.00001). (G) T12 tumor volume comparisons across treatments during 30 days of treatment. KM plot of overall survival for T12 model compared to VC n=6, MRX-2843 n=7 (HR = 0.22, p= 0.023), CTX n=6 (HR= 3.2×10⁻13, p < 0.00001), Combo n=(HR= 1.7×10⁻29, p < 0.00001 (H) Heatmap of myeloid and lymphoid population changes in 2153L tumor bearing mice across treatments. (Ly6G+) and monocytes (Ly6G-Ly6Chi) across treatments. (I) Graphical abstract for figure showing CTX reprogramming of bone marrow and monocyte differentiation differences between tumor models.

As C1q has yet to be therapeutically targeted, we focused on MERTK. We tested a MERTK kinase inhibitor (UNC2371) currently in Phase 1 clinical trials as MRX-2843 in combination with CTX. In a 30-day treatment study (Figure 2E) neither single agent produced a complete response (Figure S2C), although the chemotherapy alone was more effective. However, both models achieved a complete response (CR) (tumors were no longer palpable) with combination treatment (Figure 2F-G). CR mice were housed for 100+ days and monitored for recurrence. Long term responses (LTRs), where tumors did not recur, were achieved in 30% of 2153L tumor bearing mice (Figure 2F-G). In contrast, with treatment cessation, all T12 tumors recurred (Figure 2F–G, S2D). We confirmed the phenotype is not Balb/c specific by treating PyMT-M tumors, another mesenchymal claudin-low model in C57B/6 mice, which responded similarly to T12 (Figure S2E). To compare cancer cell direct killing efficacy of MRX-2843, we determined the IC50 values on 2153L and T12 cancer cells *in vitro.* IC50 values were similar across cancer cells (Figure S2F), and MERTK expression was restricted to TAMs and 2153L cancer cells (Figure S2G). Importantly, CTX treatment in both models skewed myelopoiesis toward monocytes and reduced neutrophils (Figure 2H), decreased immunosuppressive chemokines (Figure S2H), and drove an IFN-maintained vs. phagocytic differentiation phenotype (Figure 2I).

Overall, these data show that CTX reprograms myelopoiesis toward the monocytic lineage. However, these monocytes have different differentiation trajectories when they arrive in the microenvironments of T12 or 2153L tumors. The T12 monocytes mostly become phagocytic C1q^+^ TAMs, whereas a larger proportion of monocytes in the 2153L model maintain their monocyte profile, or differentiate into CXCL9^+^ Mo.Macs. Combining CTX with TAM receptor targeted therapy was additive, which resulted in a CR in both models, but only in the 2153L model was a LTR seen in approximately 1/3 of the animals. Compared with vehicle control, MRX-2843 (HR = 0.079, 95% CI 0.015–0.41), cyclophosphamide (HR = 0.019, 95% CI 0.0027–0.13), and the combination therapy (HR = 0.0014, 95% CI 9.4×10⁻⁵–0.022) significantly reduced the hazard of reaching humane the endpoint in the 2153L model.

### Mo.Mac antigen presentation drives differential T cell responses across TNBC subtypes

Solid tumors can significantly change host hematopoiesis resulting in a suppressive TME (33), treatment resistance (77, 78), and metastasis (79). As we observed drastic changes in the myelopoiesis with CTX treatment, we characterized the changes in the TME across the different treatments. We also investigated the PBMC populations to determine if there was any relationship with circulating immune populations with the TME. Moreover, by comparing both T12 and 2153L we correlated these cell types with the response observed in the 2153L model.

At day 18, both models responded to CTX + MRX-2843 without any apparent toxicity, and 2153L showed reduced proliferation (Figure 3A–C, S3A–B). Monocytes increased after CTX in both models, but antigen-presenting monocytes/macrophages were significantly higher in 2153L (Figure 3D-E, S3D-E). Only 2153L combination-treated tumors showed robust CD4 T cell infiltration (Figure 3F, S3C–E). By day 24, T12 tumors phenocopied the 2153L day 18 TME (Figure S3F–H). We validated that there was high CXCL9 and iNOS expression at this timepoint (Figure S3I). These data suggest monocytic differentiation supports anti-tumor immunity, with earlier CD4 recruitment distinguishing 2153L LTR potential. Next, we investigated the mechanism by which Mo.Mac reprogramming drives CD4 T cell recruitment and their function in driving LTR.

**Figure 3.**
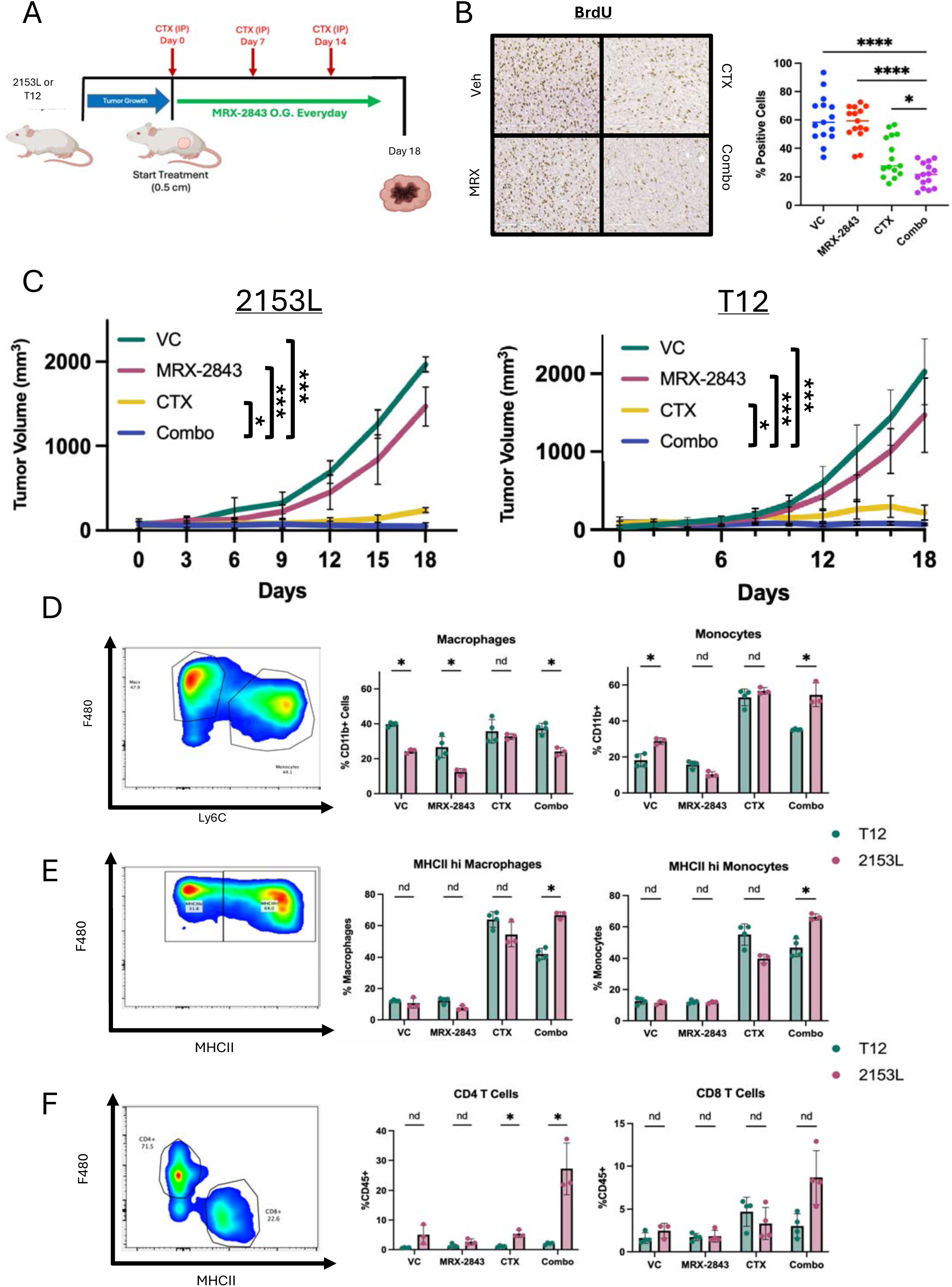
Mo.Mac antigen presentation within the TME leads to increased T cell recruitment and response in 2153L model when compared to T12. (A) Treatment strategy for day 18 analysis of TME for both 2153L and T12 murine tumors. (B) IHC staining for BrdU across treatments and quantification (Right) in 2153L model. (C) Tumor volume plot for both models across treatments for 18 days. n=5 for all groups in both models. (D) Bar plot and gating strategy for flow cytometry comparison of treatment effects on monocyte and TAM populations with the TME of both 2153L and T12 tumors. (E) Bar plot and gating strategy for flow cytometry comparison of treatment effects on MHCII expression. (F) Bar plot and gating strategy for flow cytometry comparison of treatment effects on T cell populations.

### CXCL9⁺ Mo.Macs dominate early combination-treated basal-like tumors

We performed scRNAseq on CD45^+^ cells infiltrating 2153L tumors across treatments to identify cell types, subtypes, and signaling pathways that drive response. We designed the study to interrogate temporal changes in cell populations by taking day 7 samples across all four treatments, and day 18 samples only for the CTX and Combo groups. This study design allowed for the analysis of both differential gene expression driving TME reprogramming as well as the identification of downstream consequences of cell populations and signaling.

scRNAseq across day 7 and day 18 treatments revealed T cells dominating only the day 18 combination group (Figure 4A, S4A–B). Day 7 PMNC analysis showed expansion of CXCL9^+^ Mo.Macs in CTX and combination tumors, particularly the combination group, with a concomitant decrease in C1q^+^ TAMs (Figure 4B, S4C). CXCL9^+^ Mo.Macs expressed antigen-presentation genes, IFN-response genes, and lymphocyte-activating cytokines (Figure 4C–E, S4D). CXCL9 protein was significantly increased in both models, with the highest level observed in 2153L (Figure 4H). A CXCL9^+^ Mo.Mac signature derived from these analyses correlated with TILs and associated with ICB treatment in TNBC patients (Figure 4F-G). However, the signature poorly correlated with overall patient survival within the SCAN-B, METABRIC, and CALGB chemotherapy only treated TNBC dataset (Figure S4E). These findings support an IFN-driven Mo.Mac differentiation program enhanced by MERTK inhibition (Figure 4I).

**Figure 4.**
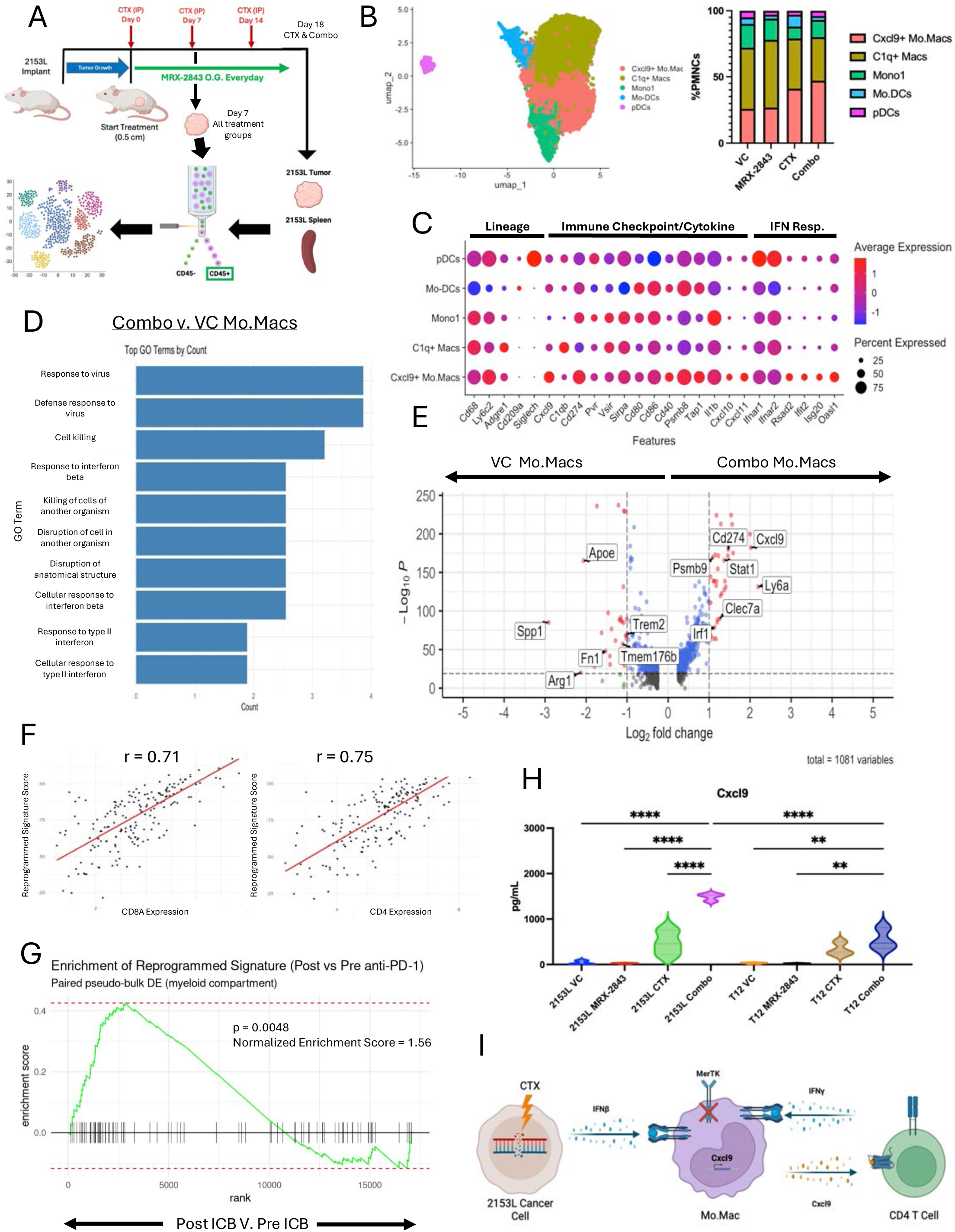
CXCL9+ Mo.Macs dominate the TME of day 7 combination treated 2153L tumors. (A) Treatment study design for scRNAseq samples at day 7 and day 18. 2153L model was used and CD45+ cells were sorted for sequencing. (B) Umap subclustering of PMNC within 2153L tumors across the four treatment arms. (C) Dot plot showing PMNC population expression of lineage, immune checkpoint/cytokines, and IFN response markers within the integrated dataset. (D) GO analysis of the top DEGs in the combination treated monocytes and macrophages. (E) Volcano plot showing top upregulated DEG in monocytes and macrophages within combination treated tumors (Right) when compared to vehicle treated monocytes and macrophages (Left). (F) Linear correlation plots showing Cxcl9+ Mo.Mac signature correlates with both CD8 and CD4 T cell infiltration in TNBC patients from the TCGA dataset. (G) GSEA of Cxcl9+ Mo.Mac signature showing signature enrichment in PD-1 treated TNBC patients. (H) Protein quantification of Cxcl9 from whole tumor lysates at day 18 across all treatments in both 2153L and T12 models. (I) Schematic showing IFN signaling synergizes with MerTKi to drive Cxcl9 expression.

Given our previous data suggesting CTX drives IFN signaling both in the bone marrow and tumors coupled with the IFN gene enrichment, we propose that this signaling drives CXCL9 expression in Mo.Macs derived from bone marrow. Additionally, the addition of MERTKi enhances the differentiation of Mo.Macs into this phenotype, resulting in increased lymphocyte recruitment and activation of the TME.

### Stem-like memory CD4⁺ T cells accumulate after combination therapy and are correlated with tumor control

With our temporal scRNAseq dataset of CD45^+^ cells across the treatments, we next extracted the lymphocyte populations and reintegrated them for subclustering analysis. Lymphocyte subclustering identified CD4, CD8, B, NK, and γδ T cells (Figure 5A). By day 18, CD4 T cells dominated combination-treated tumors, while CTX tumors were enriched for B cells (Figure 5B). CD4 T cells displayed a stem-like memory phenotype (Sell, Il7r, Bcl2) and a smaller follicular helper subset (Bcl6, Cxcr5, IL-21) (Figure S4F). CD8 effector populations were similar across treatment groups and increased when compared to VC group (Figure 5C). DEG analysis showed increased cell division and B cell activation pathways (Figure 5D), with B Cells showing increased antigen presentation and CD86 expression (Figure S5A). Cytokine profiling confirmed strong CD4-driven activation (Figure 5E). CXCR3, the receptor for CXCL9, was expressed exclusively on CD4 T cells (Figure S5B). Overall increased interactions between CD4 T cells, CXCL9+ Mo.Macs, B cells, and DCs were revealed using CellChat (Figure S4C). Lymphoid aggregates (non-lymph node, and minimally structured aggregation of T cells and B cells) were present in LTR tumor beds, while residual tumors were observed in T12 tumor bearing mice (Figures 5F). Importantly, functional studies demonstrated that blocking CD4 or CXCR3 abrogated anti-tumor activity despite continued therapy (Figure 5G–H), establishing that CXCL9/Cxcr3-dependent CD4 recruitment was essential for response.

**Figure 5.**
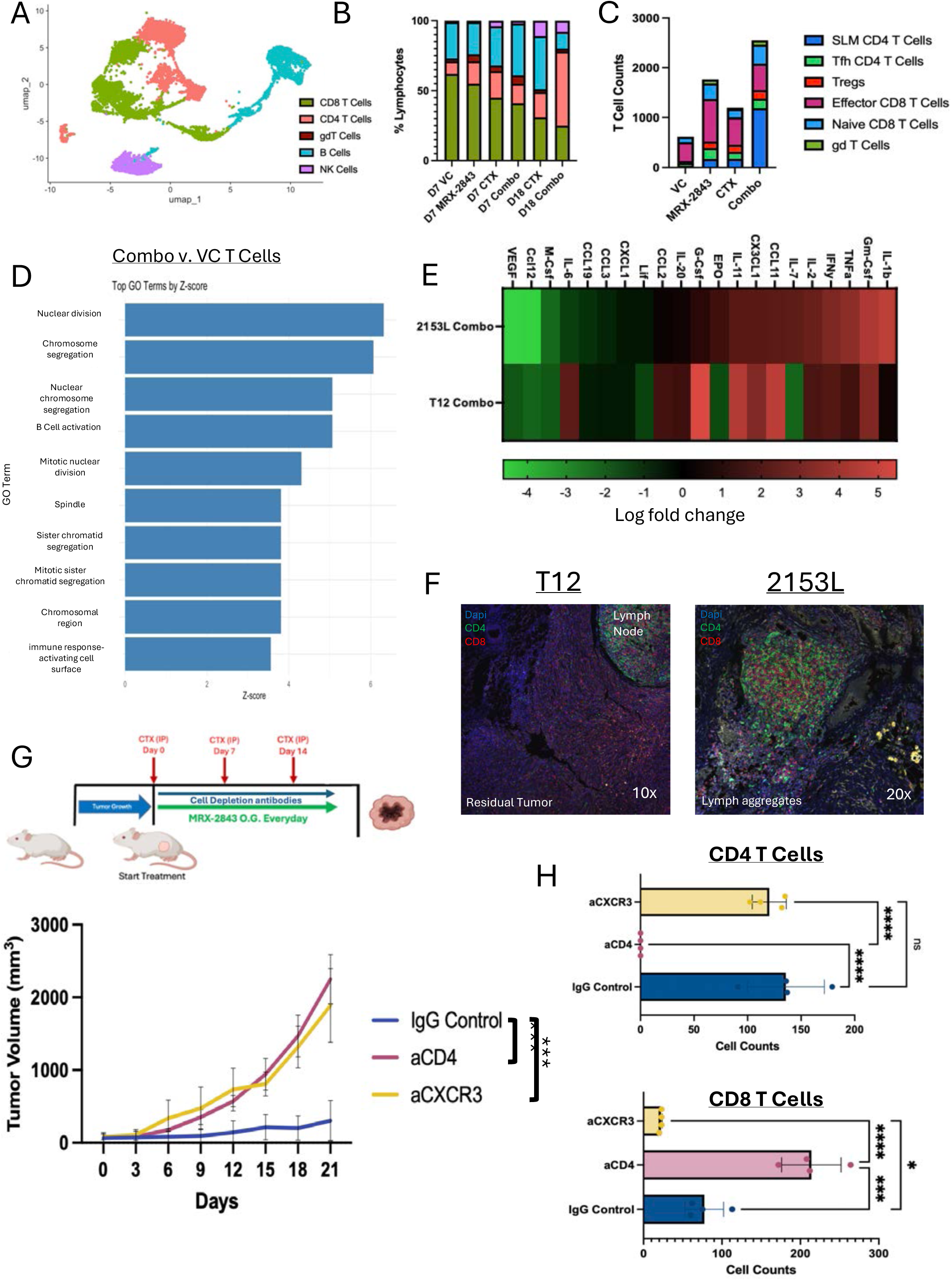
Stem-like memory CD4+ T cells dominate the TME of day 18 tumors and are necessary for anti-tumor response. (A) Umap subclustering of lymphocytes at day 7 and day 18 of treated 2153L tumors. (B) Bar plot of cell proportions across all treatments and two timepoints. (C) Total cell counts of T cell populations across all treatments and two timepoints. (D) GO analysis of top upregulated DEG in combination treated day 18 T cells when compared to day 7 vehicle treated T cells. (E) Total cytokine analysis from whole tumor lysates comparing log fold change of 2153L and T12 combination treated tumors. (F) Immunofluorescent staining of CD4 and CD8 comparing the residual fat pads of LTR 2153L mice and day 30 T12 mice. (G) Treatment strategy for CD4 depletion and Cxcr3 blockade study in 2153L model. Tumor volume plot of three different arms of study. n=4 for all groups. (H) Flow cytometry quantification of PBMC T cells populations across three different arms of study.

### TAM receptor inhibition synergizes with IFNγ/STAT1/IRF7 signaling to reprogram macrophages

We next asked what signaling pathways were altered during macrophage reprogramming. We established an in vitro system in which F4/80^+^ cells were isolated from both T12 and 2153L tumors. Once in culture, the TAMs can be treated and analyzed. For our studies we chose various approaches such as flow cytometric and western blot analyses.

*In vitro* TAM cultures showed that MRX-2843 + low-dose IFNγ induced the highest levels of CXCL9 expression and increased iNOS induction compared to other conditions (Figure 6A–B). CXCL9 protein secretion was exclusive to the combination (Figure 6C) in TAMs from both basal-like and claudin-low models (Figure 6D). Western blotting of both 2153L and T12 TAMs confirmed increased pSTAT1 and PD-L1 in the IFNγ and combination groups, decreased SOCS1 and Arg1 in the combination group, and modestly reduced p44/42 MAPK signaling with MRX-2843 (Figure 6E-F). scRNAseq data showed increased Stat1, Irf7, Irf1, and Mapk8 in CXCL9^+^ Mo.Macs (Figure 6G).

**Figure 6.**
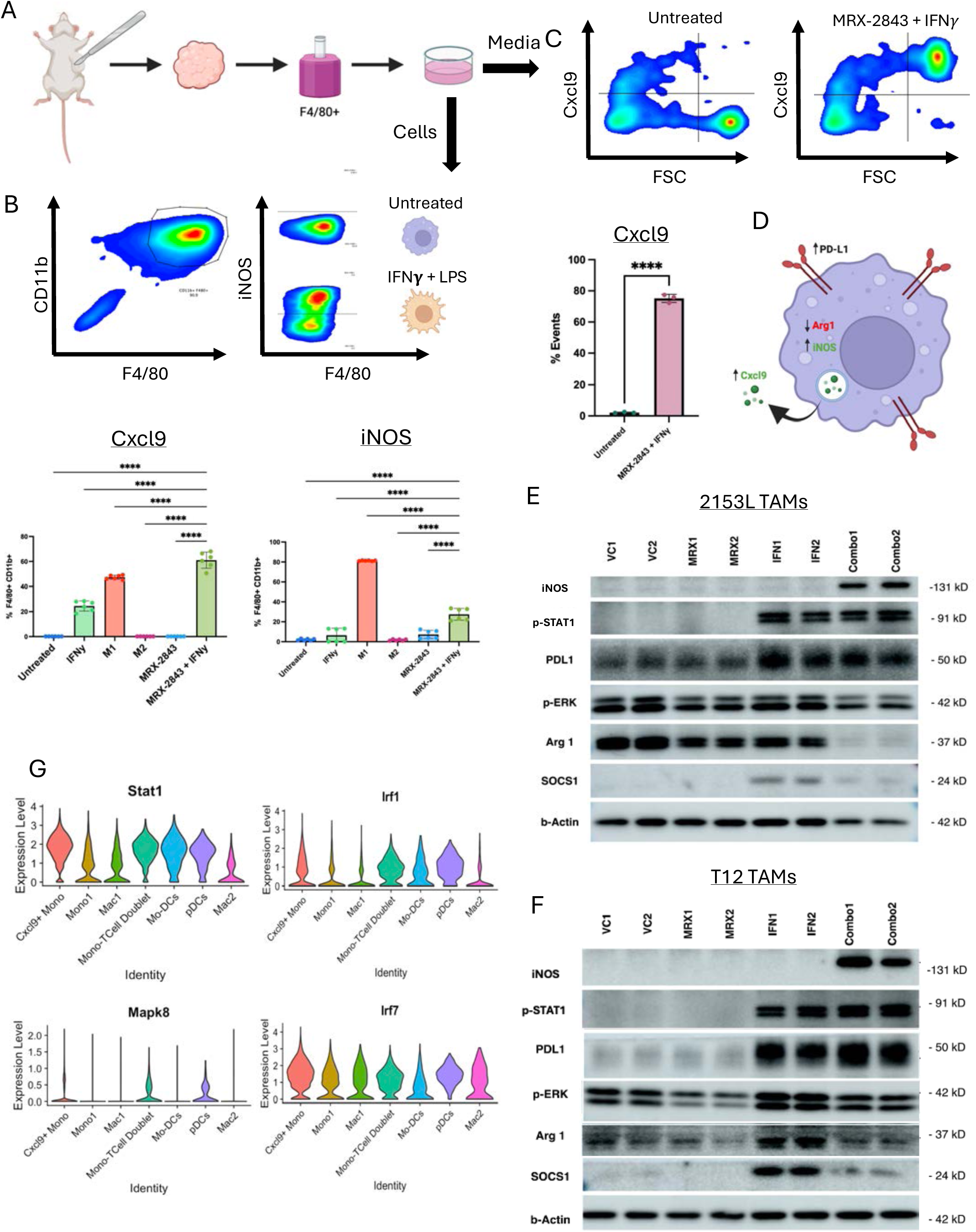
TAM receptor inhibition synergizes with IFNγ/STAT1/IRF7 signaling to reprogram macrophages. (A) Schematic of TAM extraction from murine TNBC tumors. (B) Flow cytometry analysis gating for CD11b^+^ F4/80^+^ cells after culture. iNOS can be differentially induced after “M1” reprogramming in vitro. Bar plots below summarize flow cytometry analysis across in vito treatment groups for both Cxcl9 and iNOS (C) Gating analysis comparing Cxcl9 protein differences captured in the media of MRX-2843 + IFN*γ* pulsed TAMs. Bar plot below is quantification of biological replicates. (D) Schematic of TAM expression of markers after reprogramming. (E) Western blot analysis comparing STAT, MAPK, and TAM marker expression changes across in vitro treatments in both 2153L and T12 TAMs. (G) scRNAseq of in vivo PMNCs validating in vitro pathways are similar to in vivo reprogramming pathways.

Thus, these studies demonstrate that these receptors act as a negative regulator of IFN signaling within TAMs. When TAM receptors are inhibited STAT1 signaling is hyperactivated leading to increased CXCL9 and iNOS production. Western blot analyses suggest that this regulation is correlated with MAPK downregulation and IRF1/STAT1 activation.

### PD-1 blockade enhances combination therapy and establishes durable anti-tumor immunity

Since PD-L1 expression was highest in CTX and combination tumors (Figure 7A) and localized primarily to CXCL9^+^ Mo.Macs and Mo-DCs (Figure 7B) we next asked whether addition of anti-PD-1 to this combination therapy would provide any additional benefit. Adding anti-PD-1 to CTX + MRX-2843 doubled the percentage of responding mice yielding LTRs in approximately two-thirds of animals with limited toxicity (Figure 7C–D, FigureS5D). Previously, we observed increased CD8, CD4, and B Cells within the spleens and tumor draining lymph nodes of combination treated mice (Figure S5F). In an important confirmation of the effective anti-tumor immune response, adoptive transfer of splenocytes from day 70 LTR mice significantly delayed tumor outgrowth (Figure 7E). Moreover, rechallenge of LTR mice with fresh tumor cell implantation in the contralateral mammary fat pads from both the double-combination and triple-therapy groups resulted in rejection of the newly implanted tumors (Figure 7F). Lymphoid aggregates were present in both former tumor beds (Figure 7G). B-T cell interactions in the lymph nodes suggested B cell migration and germinal center organization (Figure 7H). Overall, CXCL9^+^ Mo.Mac driven CD4 recruitment and B-cell engagement form a coordinated adaptive response (Figure 7I).

**Figure 7.**
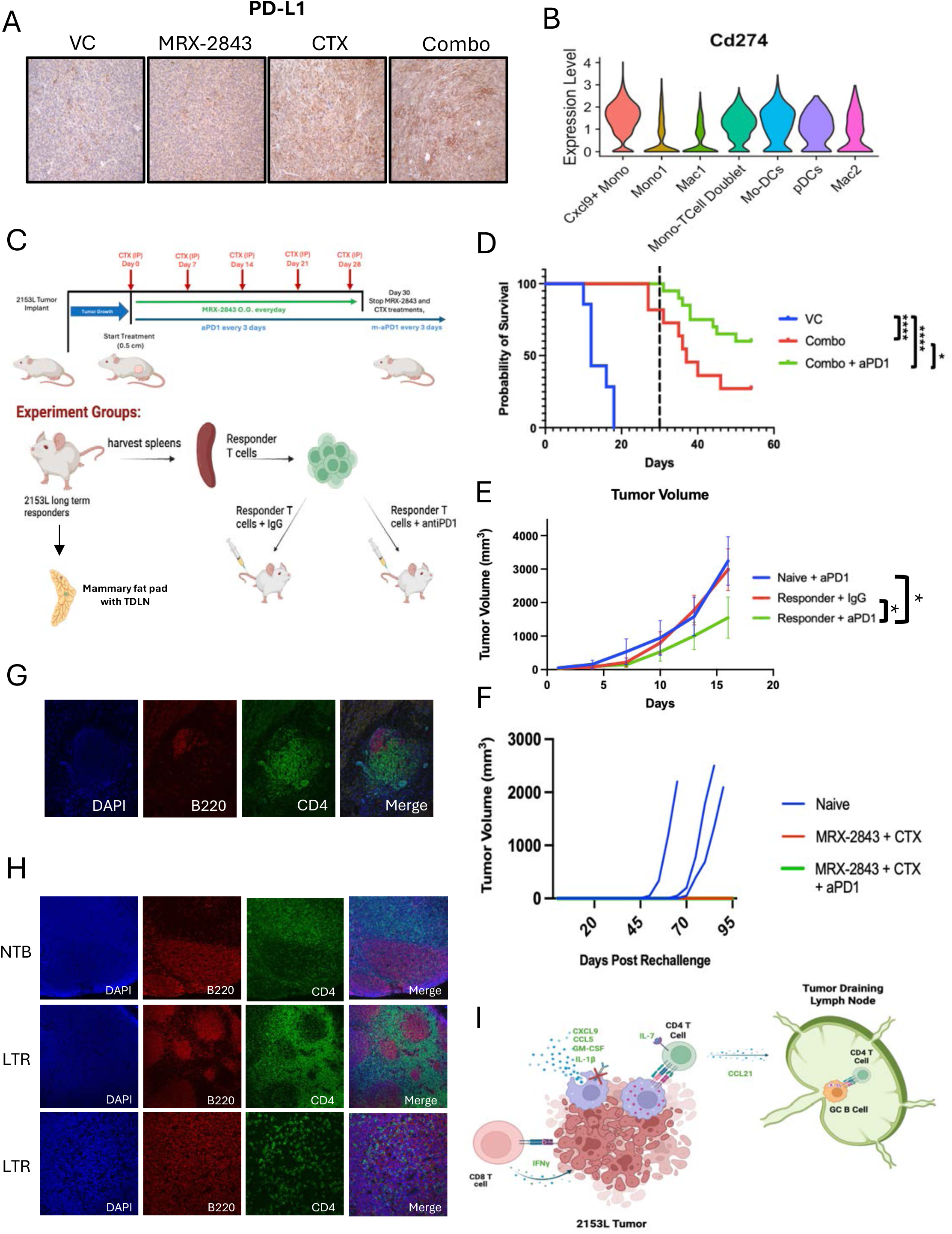
Blockade of T cell PD-1 and Cxcl9+ Mo.Mac PD-L1 interaction increases anti-tumor response of combination treatment and supports adaptive immune memory. (A) IHC staining of day 18 treated 2153L tumors. (B) Violin plot for PD-L1 expression in integrated day 7 PMNC scRNAseq dataset. (C) Treatment and sample strategy for long term treatment study of CTX + MRX-2843 + aPD1. (D) KM plot of triple combination study. VC n=7, Combo n=11, Triple Combo n=20 (E) Tumor volume plot of ACT study. n=4 for all treatment groups. (F) Tumor volume plot of rechallenge experiment. (Naïve n=3, MRX-2843 + CTX n=3, MRX-2843 + CTX + aPD1 n= 4). (G) Immunofluorescent staining of lymph aggregates containing CD20 and CD4 within the former tumor beds of LTR mice. (H) Immunofluorescent staining with CD4 and B220 of the TDLN or naïve lymph nodes within the mammary fat pad of LTR or naïve mice. (I) Summary schematic showing Cxcl9+ Mo.Macs activate and recruit CD4 T cells, which then activate B cells to drive anti-tumor memory response.

## DISCUSSION

This study establishes that C1q^+^ and CXCL9^+^ TAM populations exist in both TNBC patients and within the TME of p53null syngeneic models. We utilized p53null TNBC preclinical murine models to establish that the combination of MRX-2843 + CTX re/programs the immune response to substantially limit tumor growth. In basal-like TNBC with the greatest reprogramming of Mo Macs and infiltration of CD4 T cells, the immune response leads to LTR and anti-tumor immune memory. Finally, we establish that the addition of PD-1 to the combinatorial treatment results in most mice having a LTR in the basal-like TNBC model.

Single-cell RNA sequencing and spatial transcriptomics have revealed substantial TAM heterogeneity, with specific transcriptional states correlating with patient response to therapy (80–83). TAM populations producing lymphocyte-attracting chemokines such as CXCL9 are enriched at baseline and expand in patients responding to ICB (60–62), consistent with their role in lymphocyte recruitment and activation. In contrast, C1q^+^ macrophages correlate with poor prognosis across multiple cancer types (63–65). Although both monocytes and TAMs can present antigens and license anti-tumor T cell responses in interferon-rich contexts (84, 85), the mechanisms governing their derivation and functional polarization remain incompletely understood and appear tissue dependent. Here, we demonstrate that both CXCL9^+^ and C1q^+^ macrophage populations exist in patient samples and preclinical TNBC models, and that their relative abundance predicts the LTR response to MERTK inhibition combined with CTX and ICB. Multiple strategies have been used to therapeutically target TAMs and reverse their immunosuppressive phenotype towards an immune stimulatory function. These include HDAC inhibitors (86), pattern recognition receptor agonists (87–89), kinase inhibitors (90, 91), and engineered macrophage-based approaches (92, 93). While many showed promise preclinically, clinical translation has been limited by toxicity, insufficient efficacy, or lack of specificity. Importantly, TAM-targeting strategies alone are unlikely to induce durable anti-tumor responses, particularly in cancers such as TNBC where chemotherapy and ICB are standard-of-care. Given this therapeutic context and the heterogeneity of TNBC, evaluating TAM-modulating agents in combination regimens is critical for clinical relevance.

Cyclophosphamide is a core component of neoadjuvant chemotherapy for TNBC (69) and, at immunomodulatory doses, induces lymphodepletion followed by immune reconstitution accompanied by interferon production (70–72, 74–76). In this study, CTX reprogrammed hematopoiesis toward immune-stimulatory monocytes while suppressing neutrophil production. Although we did not longitudinally profile circulating monocytes in patients, these populations may represent clinically relevant biomarkers of response. Importantly, while neutropenia is dose-limited in patients, we observed minimal systemic toxicity in BALB/c mice treated with CTX + MRX-2843, with cachexia restricted to a subset of C57BL/6 animals.

TAM receptors play pleiotropic roles in enforcing immunosuppressive macrophage programs (25–28, 32). While MERTK-mediated efferocytosis has been extensively studied in breast cancer (30, 31), our data identify a synergistic interaction between interferon signaling and MERTK inhibition using the next-generation inhibitor MRX-2843. The drug exhibited comparable cancer cell cytotoxicity in vitro and only modestly slows growth in vivo. Together these data suggests that the interferon regulation of MERTK in Mo.Macs orchestrates the anti-tumor immune response.

Single-cell analyses revealed that CXCL9^+^ monocyte-derived macrophages constitute a minority population in untreated tumors, whereas C1q^+^ TAMs dominate both models. CXCL9^+^ Mo.Macs arise in interferon-rich niches, likely adjacent to immunogenic cell death or T cell infiltration. This supports the positive feedback of activation between IFNγ^+^ T cells and CXCL9^+^ Mo.Macs. While we were able to correlate the CXCL9^+^ Mo.Mac signature with increased T cell infiltration, the signature was poorly prognostic. This is likely attributed to the lack of single cell resolved PD-1 treated robust TNBC patient datasets. C1q^+^ TAMs express phagocytic receptors, including TREM2 and MERTK, consistent with a wound-healing, efferocytic phenotype. These pathways are likely redundant and driven by tumor-derived M-CSF (15, 94), epithelial to mesenchymal transition associated growth factors (95), and suppressive damage associated molecular pattern signaling (96). High-EMT claudin-low tumors such as T12 are dominated by C1q^+^ macrophages, potentially explaining why CSF1R inhibition, or TAM depletion, but not MERTK inhibition, induces LTRs in this context. The New START (NCT06959537) phase Ib clinical trial will test this in CL patients (40). For MERTK, we show that basal-like 2153L tumors, have higher baseline interferon signaling, which is amplified by CTX. This combined with MERTKi enables a shift of monocyte differentiation toward a CXCL9^+^ state.

Mechanistically, TAM receptor signaling restrains interferon responses through p44/42 MAPK activation and SOCS1 recruitment. CTX further induces IRF7 and IRF1 expression in CXCL9^+^ Mo.Macs, suggesting a transcriptional network in which IRF7-driven positive feedback amplifies STAT1 activity, enabling STAT1/IRF1 dependent CXCL9 transcription. Importantly, we this signaling drives TAMs into this CXCL9 phenotype in macrophages isolated from both basal-like and claudin low models, suggesting that TYRO3, AXL, MERTK family is a fundamental regulator of macrophage biology. The full efficacy of the reprogramming will depend on the context, e.g tissue of tumor origin or metastatic site and the family member complement on the infiltrating myeloid/stromal components.

Functionally, this myeloid reprogramming drives a temporally coordinated anti-tumor immune response in basal-like TNBC. Bone marrow derived CXCL9^+^ Mo.Macs recruit CXCR3+ T cells, with CD4^+^ T cells playing a non-redundant role in tumor control. Antigen-presenting Mo.Macs support expansion of stem-like memory CD4^+^ T cells, which orchestrate potential germinal center formation within tumor-associated lymphoid aggregates and tumor-draining lymph nodes. These immunological memory circuits are further amplified with the addition of PD-1 blockade. Currently, PD-L1 is a predictive biomarker in metastatic TNBC but not at the primary setting. Our data suggest that PD-L1^+^CXCL9^+^ Mo.Macs sensitize primary basal-like TNBC to PD-1 therapy. The resulting myeloid-lymphoid humoral and cellular immune memory after treatment is functionally important and sustained, as demonstrated by tumor rechallenge rejection and adoptive transfer experiments.

This study is limited by the use of pharmacologic MerTK inhibition without complementary genetic loss-of-function approache. Future studies employing conditional MerTK deletion will be required to confirm target specificity and cell-intrinsic mechanisms. In addition, direct in vivo assessment of MerTK target engagement within TAMs was not performed. Furthermore, validation across additional preclinical models will be necessary to capture the heterogeneity of human TNBC. Future studies should define the durability and functional quality of adaptive immune memory induced by this strategy, with particular emphasis on stem-like memory CD4⁺ T cells, lymphoid organization, and germinal center dynamics. Finally, biomarker-driven clinical translation of CTX plus MerTK inhibition, with or without PD-1 blockade, will be essential to determine whether this approach can expand durable immunotherapy benefit in basal-like TNBC. Taken together, our findings identify a potential therapeutically actionable axis in basal-like TNBC in which CXCL9-producing Mo.Macs link myeloid reprogramming to durable adaptive immunity. By integrating interferon-driven monocyte activation with TAM receptor blockade, CTX + MRX-2843 converts an immunosuppressive TME into one that recruits, activates, and sustains CD4^+^ T cell responses and immunologic memory. Given that basal-like tumors comprise most TNBC cases, these data provide strong rationale for translating CTX + MRX-2843 based combinations, particularly with PD-1 blockade, into clinical testing to expand durable benefit from immunotherapy.

## MATERIALS AND METHODS

### Mice

Eight- to twelve-week-old female WT Balb/c (strain047) mice were purchased from Inotiv. Eight-to twelve-week-old female WT C57BL/6J mice were purchased from The Jackson Laboratory. The mice were housed in the Transgenic Mouse Facility (TMF) animal facility at Baylor College of Medicine (BCM) in standard controlled environments. All mouse experiments adhered to the Institutional Animal Care and Use Committee (IACUC) approved AN-504.

### Tumor models

All Balb/c and *in vitro* studies utilized the 2153L and T12 models from the *Trp53*-null mammary tumor bank generated at BCM (Medina and Herschkowitz). Briefly, this tumor bank was derived via transplantation of Balb/c *Trp53-null* mammary fat pads in WT Balb/c mammary fat pads. This gave rise to a bank of heterogeneous transplantable TNBC murine tumor models, including 2153L and T12. Frozen 2153L or T12 tumor chunks were transplanted into the fourth mammary fat pads of donor WT Balb/c mice and allowed to grow to 1cm^3^. These tumors were harvested and chopped into ∼1mm^3^ chunks then transplanted into WT Balb/c mice for treatment studies. Primary cells have been generated for 2153L and T12 as previously described (ref). In vitro cancer cell studies utilized early passage cells. Both T12 and 2153L were cultured in DMEM/F-12 medium (Thermo Fisher Scientific, 11330032) supplemented with 10% fetal bovine serum (FBS) (GenDE- POT, F0900-050), 5 μg/mL insulin (Sigma-Aldrich, I-5500), 1 μg/mL hydrocortisone Sigma-Aldrich, H0888), 10 ng/mL epidermal growth factor (Sigma-Aldrich, SRP3196), and 1X Antibiotic-Antimycotic (Thermo Fisher Scientific, 15240062).

### Animal treatment protocol

Treatment studies began when tumors reached ∼62.5mm^3^. Treatment study mice were monitored daily and measurements with manual calipers and a digital scale were taken every 3 days. Mice were randomized by tumor volume and weight using the RandoMice software(ref). Mouse weight and tumor volume was measured every 3 days throughout the duration of the study. MRX-2843 (UNC2371) was kindly provided by Dr. Shelton Earp or purchased (MedChem Express, HY-101549). MRX-2843 was suspended in sterile 10% Captisol (MedChem Express, HY-17031), 10% DMSO, and PBS. CTX (Sigma-Aldrich, PHR1404-1G) was suspended in sterile PBS. MRX-2843 was administered via 200µL oral gavage at a dose of 50 mg/kg every day. 200µL of CTX was injected intraperitoneal (i.p.) once a week at a concentration of 100 mg/kg. Control mice were treated with equal volumes of both oral gavage of vehicle and i.p. injections of PBS. For the ICB study, mice were treated i.p. with anti-PD1 (BioXcell, CP151) at 200µg every three days. Control mice were injected i.p. with mouse IgG2a isotype (BioXcell, CP150) at 200µg every three days. For immune cell/signaling depletion/blockade studies mice were treated with 100 µL via i.p.

injections with either anti-CD4 (BioXCell, BE0003-1), anti-CXCR3 (BioXCell, BE0249), or control Armenian hamster IgG (BioXCell, BE0091) and rat IgG2b (BioXCell, BE0090). The anti-CD4 group was given 400µg of antibody per dose, while anti-CXCR3 was given 200µg per treatment. BioXCell antibodies were diluted with InVivoPure Dilution Buffer (BioXCell, IP0070).

### Adoptive T cell transfer and cancer cell rechallenge

Splenocytes were harvested from LTR mice that had been treated with the combination of MRX-2843 + CTX + aPD1 or from WT non-treated Balb/c mice. Spleens were minced, then passed through a 40µm filter. This was followed by a brief RBC lysis (BioLegend, 420301). Both CD4+ and CD8+ T Cells were negatively selected using the EasySep Mouse T Cell Isolation Kit (STEMCELL Technologies, 19851) and EasySep Magnet (STEMCELL Technologies, 18000). T Cells were counted and resuspended in PBS, then 1 million cells were injected via tail vein into each 2153L tumor bearing mice. Mice were simultaneously treated with either anti-PD1 (BioXCell, CP151) or IgG2a control (BioXCell, CP150). LTR mice from both the double and triple combination groups or treatment naïve WT Balb/c mice were rechallenged in the contralateral mammary fat pad with 10,000 2153L cells freshly isolated from a 2153L primary tumor bearing mouse (See TAM extraction protocol below).

### TAM isolation and *in vitro* treatments

TAMs were isolated directly from both T12 and 2153L mammary tumors by using either CD11b+ selection (STEMCELL Technologies, 18970) or F4/80+ selection (STEMCELL Technologies, 100-0659) utilizing the EasySep Magnet (STEMCELL Technologies, 18000). Tumors were harvested, minced into a paste, and digested in collagenase A (Sigma-Aldrich, 11088793001) for 1.5hrs at 37°C with 125rpm rotation followed by a brief centrifugation to pellet for tumor organoids/cells, leaving stromal/immune cells enriched in the supernatant. Stromal cells were then pelleted, counted, and diluted according to manufacturer protocol for positive selection. After selection, TAMs were counted and seeding was dependent on well plates used: low-attachment 24 well plates (300,000 cells), tissue culture treated 24 well plates (250,000 cells), tissue culture treated 96 well plates (35,000 cells), and 8-well chamber slides (80,000 cells, Corning, 354108). TAMs were cultured in RPMI-1640 media (GenDEPOT, CM059-050) supplemented with 20% FBS, 100µM sodium pyruvate (Thermo Fisher Scientific, 1130032), MEM Non-Essential Amino Acids Solution (Thermo Fisher Scientific, 11140050), Pen/Strep Antibiotics (GenDEPOT, CA002010), 55µM ß-Mercaptoethanol (Thermo Fisher Scientific, 21-985-023), and 10ng/mL mouse M-CSF (Biolegend, 576406). TAMs were allowed to adhere to plates O/N, then washed with PBS to remove cell debris. This was then followed with the addition of media with diluted drug treatments. The treatment groups included: DMSO untreated, 1µg/mL LPS (Millipore Sigma, L2880) + 50ng/mL IFNγ (BioLegend, 575306), 10ng/mL IL-4 (BioLegend 574304), 10ng/mL IL-13 (Fisher Scientific, 413ML005), 2µM MRX-2843, or MRX-2843 + IFNγ. TAMs were incubated for 24 hours in treatment media. The following day the TAMs are retreated with fresh reagents. For intracellular secreted factor interrogation (CXCL9), TAMs were treated with Brefeldin A (BD Bioscience, 555029) to trap the cytokine inside the cells during second treatment. Six hrs after retreatment TAMs were lysed for protein with RIPA Buffer (Thermo Fisher Scientific, P189900) with 1X protease and phosphatase inhibitors, lysed for RNA with TRIzol reagent (Thermo Scientific, 15596018), or collected for flow cytometry.

### Immunoblotting

Once TAMs were lysed, protein was quantified with a BCA assay (Thermo Fisher Scientific, 23225). Lysates were diluted into loading buffer (BioRad, 1610747 and ßME), then heated to 95°C for 10min to denature proteins. Proteins were equally loaded, electrophoresed on SDS-PAGE gels and then transferred to a PVDF membrane (Millipore, IPVH00010). Blots were blocked with 5% non-fat dry milk (BioRad, 1706404) in PBS, then probed with the following antibodies in 5% BSA solution: MERTK (Cell Signaling Technology (CST), 38102), AXL (CST, 8661), p44/42 (CST, 9102), p-p44/42 (CST, 9101), iNOS (CST, 13120), PD-L1 (R&D Systems, AF1019), p-STAT1 (CST, 8062), Arg1(CST, 93668), and SOCS1 (CST, 55313). Secondary antibodies were diluted in 5% non-fat milk and incubated on shaker at RT for 1hr. Blots were incubated in chemiluminescence substrate buffer (Thermo Scientific, 32106), then imaged.

### Immunohistochemistry and immunofluorescence

Primary tumors, mammary glands with TDLNs, and spleens were submerged and fixed with 4% paraformaldehyde for 24h at 4°C, then transferred to 70% ethanol. Samples were then paraffin embedded and sectioned into 5um sections. SDS-PAGE gel electrophoresis H&E samples were deparaffinized and rehydrated then subsequently stained with Harris Hematoxylin (Poly Scientific, S176) and Eosin (Fisher Scientific, 22-050-198). IHC samples were deparaffinized and rehydrated followed by antigen retrieval with either citrate buffer (pH 6.0) or Tris-EDTA buffer (pH 9.0) for 30 min in a steamer. Endogenous peroxidase was blocked with 3% H2O2 for 10 min. Samples were then incubated with 5% goat serum in PBS blocking buffer for 1hr at room temperature. Primary antibodies were diluted in blocking buffer and incubated at 4°C O/N. The following primary antibodies and dilutions were used: F4/80 (CST, 70076 1:1,000), CD8α (CST, 98941, 1:500), CD4 (Abcam, ab183685, 1:500), BrdU (Abcam, ab6326, 1:1,000), and Vimentin (CST, 5741, 1:1,000). The anti-rabbit (Vector Laboratories, PI-1000-1) biotin conjugated secondary antibody was incubated at room temperature RT for 1hr. ABC reagent (Vector Laboratories, PK7100) followed by DAP peroxidase substrate (Vector Laboratories, sk-4105) were used to develop staining. Hematoxylin (see above) counterstains were applied, and slides were mounted with cover slips and allowed to dry O/N. Both H&E and IHC slides were imaged using the Olympus BX40 light microscope and MPX-5C pro low-light camera. Slides were also scanned with the Aperio ImageScope (Leica Biosystems) allowing for quantification using the Aperio ImageScope software (v12.3.3.5048). At least three different samples and three representative images were utilized for quantification.

FFPE Immunofluorescence samples were deparaffinized, rehydrated, antigen retrieved, and blocked with 5% goat serum similarly to above. Primary antibodies were diluted in PBS blocking buffer and incubated at 4°C O/N. The following primary antibodies and dilutions were used: CD4 (Abcam, ab183685, 1:500), CD20 (CST, 70168, 1:500), CD8a (CST, 98941, 1:500), CD8a (eBioscience, 14-0081-82, 1:500), CD45R (BD Biosciences, 550286, 1:500), and Ly6C (Abcam, ab314120, 1:100). Appropriate fluorophore conjugated secondary antibodies (Alexa Fluor, Invitrogen) were incubated for 1hr at RT. NucBlue (Invitrogen, R37606) nuclear stain was diluted in PBS and incubated for 15min at RT. Slides were mounted with Aqua/Poly-mount (Fisher Scientific, NC9439247) and allowed to dry in 4°C O/N. Samples were imaged with a Nikon A1-Rs confocal microscope at 10x, 20x, and 40x magnifications. Tif files were processed and converted with the Imagj software.

### Flow cytometry and cell sorting

Tumors were processed, digested, and immune cells were enriched as previously described above. Stromal/immune pellets were resuspended in RBC lysis buffer for 5 min at RT. Cells were then passed through a 40µm cell filter. Cells were counted, spun, then resuspended to 10 million cells/mL in FACS buffer (PBS +2% FBS). Two million cells/sample were used for staining and analysis. Cells were stained with Live/Dead Fixable Yellow (Thermo Fisher Scientific, L34968, 1:800) diluted in PBS, then blocked with anti-mouse CD16/CD32 antibody (BioLegend, 101320), followed by extracellular antibody staining on ice for 1hr. Cells were fixed and permeabilized with the FOXP3/Transcription Factor Staining Buffer Kit (Thermo Fisher Scientific, 005523-00) O/N. The following morning the cells were blocked with 2% rat serum diluted in 1X permeabilization buffer and incubated with intracellular antibodies The following antibodies and dilutions were used for tumor infiltrating immune cells identification: CD45 (BioLegend, 103128, 1:250), MHCII (BioLegend, 116513, 1:500), CD11b (BioLegend, 101228, 1:400), Arg1 (eBiosceinece, 12-3697-82, 1:200), CD206 (BioLegend, 141732, 1:150), Ly6C (BioLegend,128017, 1:300), F4/80 (BioLegend,123116, 1:150), Ly6G (BioLegend, 127652, 1:200), iNOS (), CD45 (Biolegend, 103126, 1:300), CD3 (BioLegend, 100328, 1:100), Foxp3 (BioLegend,126404, 1:150), CD45R (BioLegend, 103222, 1:200), Granzyme B (Biolegend, 372204, 1:200), CD8 (BioLegend, 100730,1:200), and CD4 (BioLegend, 100460,1:100). Compensation beads (Thermo Scientific, 01-2222-42) were used for single color controls and compensation. Monocytes are defined as CD45^+^CD11b^+^Ly6C^hi^Ly6G^−^F4/80^−^. Macrophages are defined as CD45^+^CD11b^+^Ly6C^−^Ly6G^−^F4/80+. Neutrophils are defined as CD45^+^CD11b^+^Ly6C^hi^Ly6G^−^F4/80^−^. Cytotoxic T cells are defined as CD45^+^CD3^+^CD8^+^CD4^−^. Helper T cells are defined as CD45^+^CD3^+^CD8^−^CD4^+^.

Blood was obtained via retro-orbital bleeds prior to euthanasia. 30µL of blood was diluted in 1X RBC lysis buffer for 30min. Bone marrow was collected from femurs of hind legs. These cells were directly filtered through a 40µm filter and stained via the same protocol above. Cells were then stained by following the same protocol above. The only deviation being dapi (Invitrogen, R37606) utilization for live/dead gating. The following antibodies and dilutions were used for mature immune cell identification: CD45 (BioLegend, 103128, 1:500), CD11b (BioLegend, 101227, 1:400), Ly6C (BioLegend, 128017, 1:300), Ly6G (BioLegend, 127651, 1:200), CD3, CD45R, CD8, and CD4 (see above for vendor).

TAMs were lifted from low attachment 24 well plates after two days of culturing and treatments with PBS manual washes. TAMs were transferred to U-bottom 96 well plates for staining. TAM marker staining followed protocol mentioned above. The following antibodies and dilutions were used for TAM programming interrogation: CD11b (BioLegend, 101227, 1:500), F4/80(BioLegend, 123116, 1:150), CXCL9 (BioLegend, 515604, 1:200), iNOS (Miltenyi Biotec, 130-116-421, 1:100), MHCII (BioLegend, 107643, 1:500), Arginase1 (BioLegend, 165803, 1:200), and CD206 (BioLegend, 141732, 1:150). Media was harvested after two days of culture across treatment groups. Media was processed via mouse BioLegend LEGENDplex assay V-bottom plate procedure (BioLegend, 741024).

Tumor, TAMs, blood, and bone marrow samples were processed on the Attune NxT cytometer within the Cytometry and Cell Sorting Core at Baylor College of Medicine. Event data was processed, compensated, and gated with the Flowjo v10 software.

### scRNAseq

Day 7 mice were treated 3 hours before euthanasia and harvesting. Tumors were processed, digested, and diluted as mentioned above. Spleens were harvested as mentioned above. Anti-CD16/32 antibody was used to block FcR binding, then incubated with anti-CD45 (BioLegend, 103128 1:250) for 45 min. Cells were then stained in collection buffer (PBS + 20%FBS) with Dapi. Minimally, 150,000 cells were sorted for Dapi-CD45+ on a Sony MA900 in the Cytometry and Cell Sorting Core at Baylor College of Medicine. After sorting, cells were spun, resuspended in PBS, and submitted to the Single Cell Genomics Core at Baylor College of Medicine.

A single cell 3’ Gene Expression Library was prepared according to Chromium Single Cell Gene expression 3’V4 On-Chip Multiplexing kit (10x Genomics). In brief, single cells, reverse transcription (RT) reagents, Gel Beads containing barcoded oligonucleotides, and oil were loaded on a Chromium X (10x Genomics) to generate single cell GEMS (Gel Beads-In-Emulsions) where full length cDNA was synthesized and barcoded for each single cell. Subsequently the GEMS are broken and cDNA from each single cell is pooled. Following cleanup using Dynabeads MyOne Silane Beads, cDNA is amplified by PCR. The amplified cDNA is fragmented to optimal size and the 3’ Gene Expression (GEX) library was generated via End-repair, A-tailing, Adaptor ligation and PCR amplification. Libraries were sent to the Genomic and RNA Profiling Core for sequencing with a NextSeq 500. Raw sequencing data were processed using Cell Ranger v9.0.1 (10x Genomics) after receiving the raw output files from the Genomic and RNA Profiling Core (GARP) at Baylor College of Medicine. A configuration file was used for sample demultiplexing, containing the correspondence between OCM Barcodes (OB) and sample names (OB1: *Combination_CD45*; OB2: *CTX_CD45*; OB3: *MRX_CD45*; OB4: *VC_CD45*). The Cell Ranger pipeline generated count matrices and cell barcode files separately for each sample.

Downstream analyses were performed using Seurat v5.2.0 (97). Single-cell objects were created and filtered based on standard quality-control criteria (minimum features per cell: 250–6,000; mitochondrial gene fraction < 15%). Datasets were normalized then, to correct for batch effects, we applied Seurat’s *FindIntegrationAnchors* and *IntegrateData* functions prior to generating the final integrated single-cell objects. Standard neighborhood and clustering analyses from Seurat pipeline were used to generate clusters at 0.5 resolution. Clusters were first unbiasedly identified with SingleR Immgen (https://github.com/SingleR-inc/celldex), then verified manually with lineage markers. Differential gene expression was identified with the FindMarkers function from the Seurat package, which used the MAST model testing between two clusters or datasets. Other packages used for downstream analyses include CellChat, UCell, Slingshot, and EnhancedVolcano.

### Cytokine Profiling

Snap-frozen tumor chunks for treatment studies were homogenized with bead homogenizer (BeadBlaster 24) in T-PER buffer with protease inhibitor cocktail. Protein concentration was determined with a BCA assay. Tumor homogenates were diluted to 1.8 µg/µL. Plasma was isolated by spinning fresh blood from treated mice in EDTA lined tubes. Plasma was isolated and diluted 2-fold with PBS. All samples were mailed to Eve Technologies Corp. (Calgary, Alberta, Canada) on dry ice. All samples were profiled using the 44-Plex mouse discovery panel.

### Statistics

See figure legends for animal numbers, replicates, and specific statistics used for each experiment. For comparisons of 2 groups, an unpaired Student’s t test was used. For comparisons of 3 or more groups and pairwise comparisons, an ordinary one-way ANOVA was used with Tukey’s multiple -comparison test. For mouse tumor volume and body weight analyses, a two0way ANOVA with Šidák’s multiple comparison test was used. For Kaplan-Meier survival analyses, a log-rank test for multiple comparisons with Bonferroni’s method was used. Survival was analyzed using a Cox proportional hazards model with humane endpoint as the event and right-censoring at the fixed study endpoint (day 100). A P value of less than 0.05 was considered significant. Statistical analyses were mostly conducted with GraphPad Prism 10 or Microsoft Excel. scRNAseq statistical analyses were conducted in R.

### Human and mouse comparison

Single-cell RNA sequencing data from a cohort of human breast cancers was downloaded from the Gene Expression Omnibus (GSE176078). Single-cell objects were created and filtered based minimum features per cell and mitochondrial gene fraction 10%. Data was then filtered to contain only samples from triple-negative breast cancer. Datasets were log-normalized and clustered using the standard Seurat pipeline described above. Clusters were labeled for cell type by SingleR and lineage markers as described above. Datasets were then filtered to exclude non-myeloid cells, and normalization and clustering pipelines were repeated on these myeloid populations at 0.2 resolution. Clusters were re-labeled for cell type by SingleR and lineage markers as described above. Marker genes defining clusters were identified using the FindAllMarkers function from the Seurat package. Signatures for cell types were derived by taking the top 50 differentially expressed genes for each cluster. Clusters matching expression patterns of CXCL9+ and C1q+ macrophages were identified in these human samples. Differential gene expression between these populations performed using the FindMarkers function, and Gene Set Enrichment Analysis (GSEA) was performed to identify differentially expressed pathways using the fgsea (v1.32.4) package in R and Gene Ontology:Biological Processes pathway set from the Molecular Signatures Database (MSigDB).

Single cell RNA-sequencing data from human breast cancer samples before and after anti-PD-1 therapy was downloaded from a public repository hosted by the VIB KU Leuven Center for Cancer Biology (https://lambrechtslab.sites.vib.be/en/single-cell). Single-cell objects were created and filtered based minimum features per cell and mitochondrial gene fraction 10%. Datasets were normalized, clustered, and annotated for cell type as above. A reprogrammed macrophage signature was derived from the murine single-cell RNA-sequencing data as above using the top 100 differentially upregulated genes from day 7 combination treated Mo.Macs scRNAseq dataset and converted to human orthologs. Macrophages and monocytes were pseudobulked using the AggregateExpression function in Seurat grouped by patient ID and treatment timepoint (pre- and post-therapy). DESeq2 (v1.46.0) was used to model gene-level counts with a paired design for differential gene expression between pre- and post-treatment samples. GSEA was then performed between pre- and post-treatment timepoints for the reprogrammed macrophage signature.

Bulk RNA sequencing and clinical data for the TCGA breast cancer cohort (TCGA-BRCA) were downloaded from cBioPortal. PAM50 subtype calls were taken from Thennavan, et al. (https://www.ncbi.nlm.nih.gov/pmc/articles/PMC9028992/). Samples were scored for CXCL9+ and C1Q+ macrophage signatures by single-sample Gene Set Enrichment Analysis (ssGSEA) using the GSVA package (v2.0.7) in R. Basal and claudin-low samples were binned into CXCL9 high, C1Q low and CXCL9 low, C1Q high groups based on median scores of CXCL9+ and C1Q+ macrophage signatures. Kaplan-Meier survival analysis was performed on corresponding patients for these samples using the survival (v3.8.3) and survminer (v0.5.0) packages in R.

## Data Availability

scRNAseq raw and processed data has been deposited in the NCBI GEO database (accession number: GSE318315). CXCL9/C1Q signatures are provided in supplementary tables 1-2. All reagents developed in this study are available from the corresponding or lead contact following a completed materials transfer agreement. Please direct information requests to the corresponding or lead contacts.

## Author Contributions

A.J.S. conducted experiments, analyzed data, and wrote the manuscript. A.J.S., D.A.P., and N.Z. conceptualized mouse study design and experiments. A.J.S. and Z.S. conceptualized human and mouse bioinformatic comparisons. Z.S. conducted the analysis for human datasets. N.G., S.J.C., X.Y., and Z.G. assisted in conducting experiments and processing samples. Y.G., F.L., and C.R. shared scRNAseq datasets and assisted in processing data. S.H. and ZC.S. designed RPPA assay and assisted in RPPA data processing. S.E. and C.M.P. shared drug compound, conceptualized study designs and data interpretations, and edited the manuscript. J.M.R. conceived, supervised, supported all studies, as well as edited the manuscript.

## Acknowledgments

We are grateful for the mentorship, critical feedback, and support from the Rosen laboratory members and our co-authors Drs. Shelley Earp and Chuck Perou. We thank Drs. Xiang Zhang, Valentina Hoyos, David Rowley, and William Decker for their feedback and support while serving on A.J.S. thesis advisory committee. We thank Drs. Fenglue Peng, Yang Gao, and Swarnima Singh for sharing the cell/feature matrix files for integration analysis. This project was supported by the Cytometry and Cell Sorting Core at Baylor College of Medicine with funding from the CPRIT Core Facility Support Award (CPRIT-RP240432) and the NIH (CA125123, OD036336, and OD038251), and the assistance of Joel M. Sederstrom, the Single Cell Genomics Core at Baylor College of Medicine with funding from the CPRIT RP250580 and NIH P30CA125123, the Integrated Microscopy core supported by NIH (DK56338, CA125123, ES030285), and CPRIT (RP150578, RP170719), and the Pathology Core of Lester and Sue Smith Breast Center at BCM. A.J.S. is supported by Susan G Komen (**ASP241264627,** SAC232150). Z.S. is supported by funding from the NIH (UNC-Integrated Translational Oncology T32-CA244125 to UNC/ZS). D.A.P. is supported by the American Cancer Society Postdoctoral Fellowship (PF-22-163-01-MM) and NIGMS MOSAIC (1K99GM155594-01). N.Z. is supported by Cancer Prevention and Research Institute of Texas (CPRIT) (RP220468). J.M.R., C.M.P. and N.G. are supported by NIH (R01CA148761-15) and Susan G Komen (SAC232150). The work received support for the Breast Cancer Research Foundation to Drs. Earp and Perou. C.M.P. was supported by NCI Breast SPORE program (P50-CA058223) and Breast Cancer Research Foundation (BCRF-23-127). All illustrations were created with Bio Render.

