## Supplemental Figures for "MERTK inhibition cooperates with immunomodulatory cyclophosphamide to induce CXCL9⁺ monocyte-macrophage programming and durable anti-tumor immunity in triple negative breast cancer"

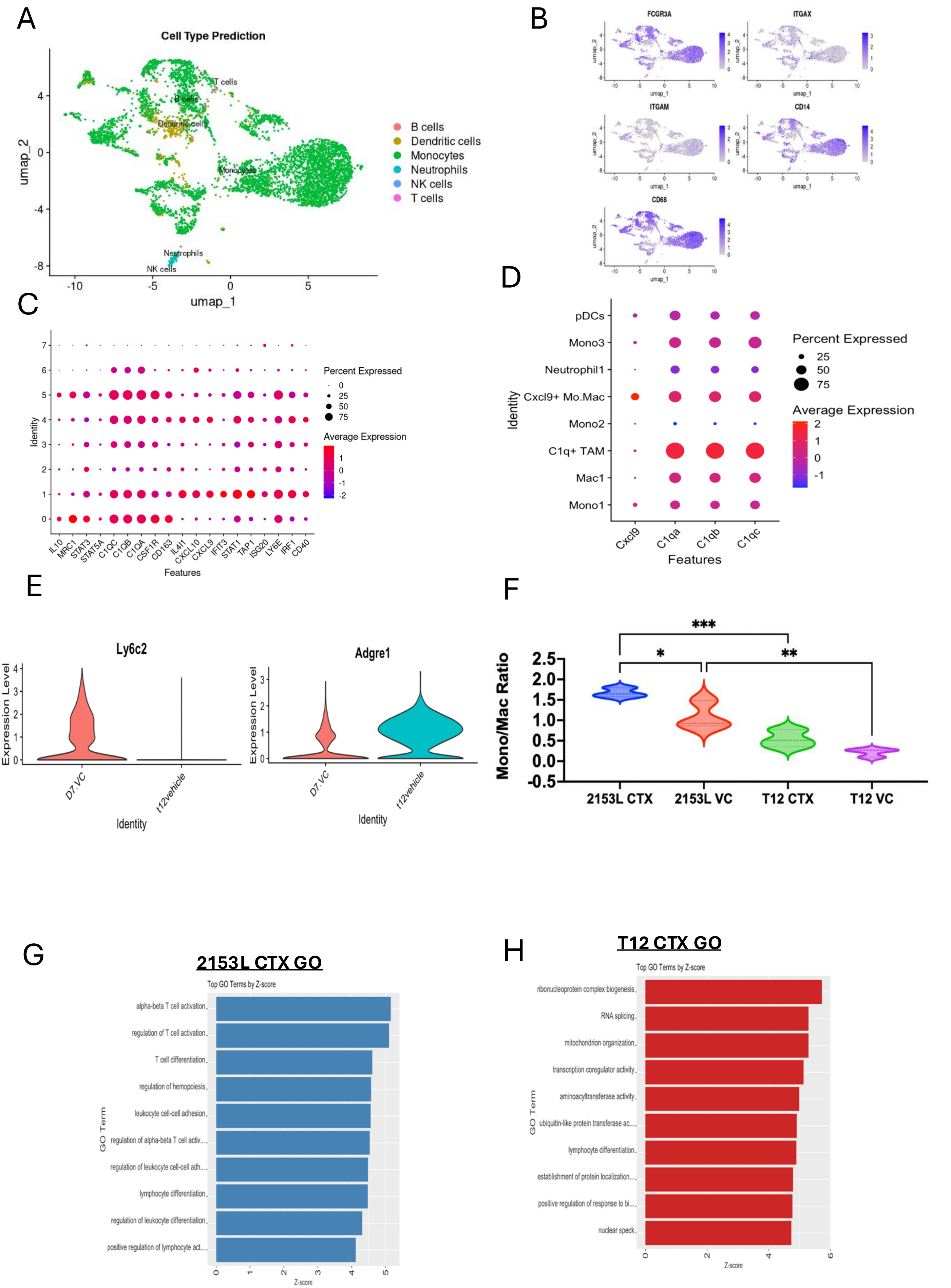
A
Cell Type Prediction
B cells
Dendritic cells
Monocytes
Neutrophils
NK cells
T cells
umap_1
umap_2
B
FCGR3A
ITGAX
ITGAM
CD14
CD68
C
Percent Expressed
Average Expression
Identity
Features
IL10 MRC1 STAT3 STAT5A C1QC C1QB C1QA CSF1R CD163 IL4I1 CXCL10 CXCL9 IFIT3 STAT1 TAP1 ISG20 LY6E IRF1 CD40
D
pDCs
Mono3
Neutrophil1
Cxcl9+ Mo.Mac
Mono2
C1q+ TAM
Mac1
Mono1
Cxcl9 C1qa C1qb C1qc
Features
E
Ly6c2
Adgre1
Expression Level
D7.VC
t12vehicle
Identity
F
Mono/Mac Ratio
2153L CTX
2153L VC
T12 CTX
T12 VC
G
2153L CTX GO
Top GO Terms by Z-score
alpha-beta T cell activation
regulation of T cell activation
T cell differentiation
regulation of hemopoiesis
leukocyte cell-cell adhesion
regulation of alpha-beta T cell activ...
regulation of leukocyte cell-cell adh...
lymphocyte differentiation
regulation of leukocyte differentiation
positive regulation of lymphocyte act...
GO Term
Z-score
H
T12 CTX GO
Top GO Terms by Z-score
ribonucleoprotein complex biogenesis
RNA splicing
mitochondrion organization
transcription coregulator activity
aminoacyltransferase activity
ubiquitin-like protein transferase ac...
lymphocyte differentiation
establishment of protein localization...
positive regulation of response to bi...
nuclear speck
GO Term
Z-score

**Supplemental Figure 1. p53null murine TNBC tumors model PMNC populations in patients.** (A) UMAP of PMNC populations. (B) Feature plots showing RNA expression of lineage markers in the various clusters. (C) Dot plot expression of immune regulatory genes including Cxcl9 and C1q in human dataset. (D) Dot plot of Cxcl9 and C1q expression in PMNCs from p53null murine models. (E) Lineage marker expression comparing T12 to 2153L. (F) Flow cytometry analysis of Mono/Mac ratio ($LY6C^+/GR1^-F4/80^+$). (G) GO analysis comparing total mono/mac populations in 2153L CTX treated group to 2153L VC treated group. (H) GO analysis comparing total mono/mac populations in T12 CTX treated group to T12 VC treated group.

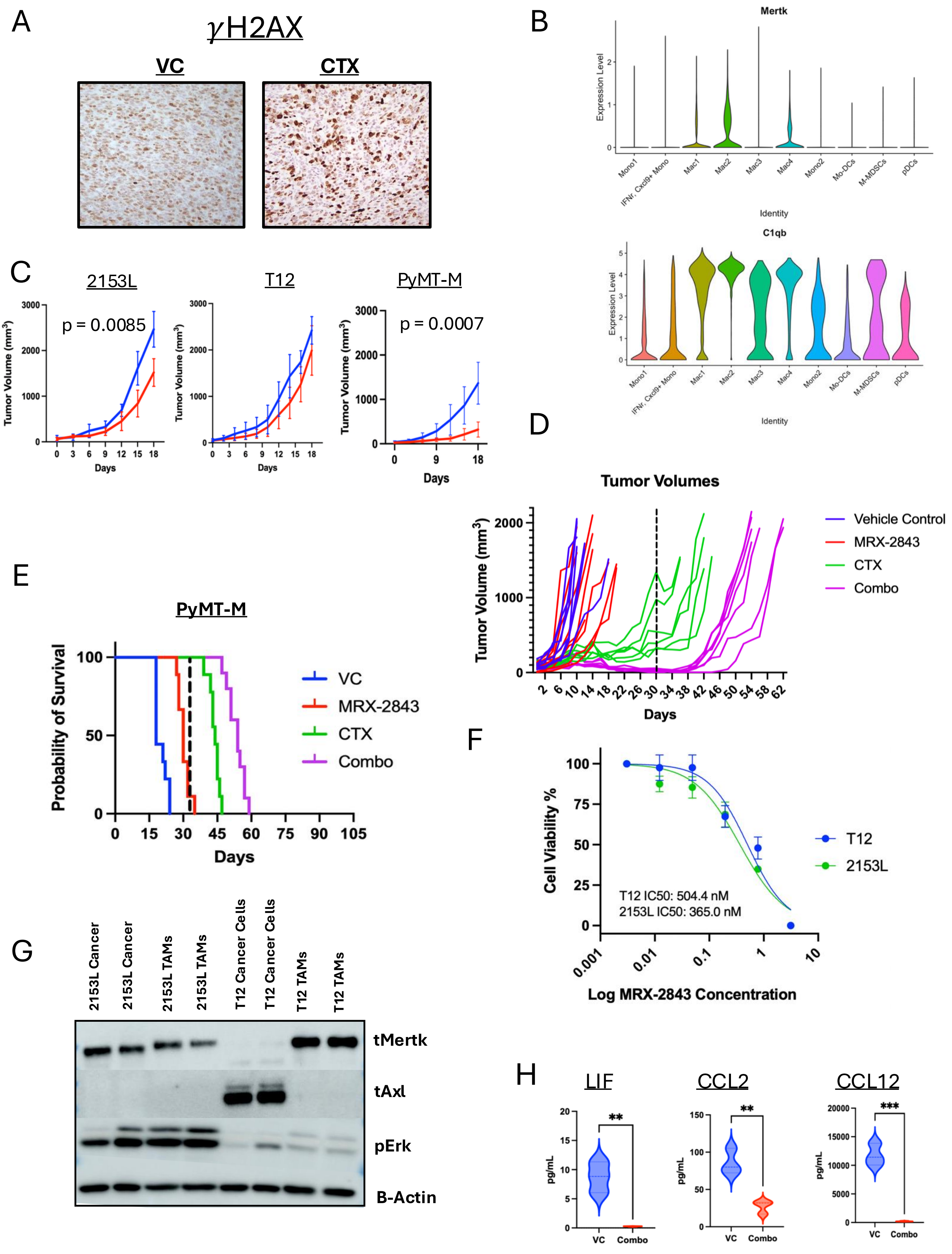
A
γH2AX
VC
CTX
B
Mertk
Expression Level
Identity
C1qb
Mono1
IFNr, Cxcl9+ Mono
Mac1
Mac2
Mac3
Mac4
Mono2
Mo-DCs
M-MDSCs
pDCs
C
2153L
T12
PyMT-M
p = 0.0085
p = 0.0007
Tumor Volume (mm³)
Days
D
Tumor Volumes
Vehicle Control
MRX-2843
CTX
Combo
E
PyMT-M
Probability of Survival
VC
MRX-2843
CTX
Combo
Days
F
Cell Viability %
T12
2153L
T12 IC50: 504.4 nM
2153L IC50: 365.0 nM
Log MRX-2843 Concentration
G
2153L Cancer
2153L Cancer
2153L TAMs
2153L TAMs
T12 Cancer Cells
T12 Cancer Cells
T12 TAMs
T12 TAMs
tMertk
tAxl
pErk
B-Actin
H
LIF
CCL2
CCL12
pg/mL
VC
Combo
**
**
***

**Supplemental Figure 2. MerTK is a viable target in p53null models leading to CR in p53null tumors when treated with MRX-2843 + CTX.** (A) $\gamma$H2AX staining showing increased DNA damage in CTX treated 2153L tumors. (B) MERTK and C1Q are co-expressed in suppressive TAM populations. (C) MRX-2843 modestly slows tumor growth in 2/3 models. (D) Tumor volume plot of T12 treatment study showing recurrence after treatment cessation. (E) KM plot of PyMT-M tumor bearing C57BL/6 mice treated with MRX-2843 + CTX. (F) IC50 curves showing similar cancer cell killing profile in both p53null models in vitro. (G) Western blot analysis showing differential TAM receptor and MAPK expression. (H) Protein analysis of immunosuppressive chemokine expression in VC or Combo treated 2153L tumors at day 18.

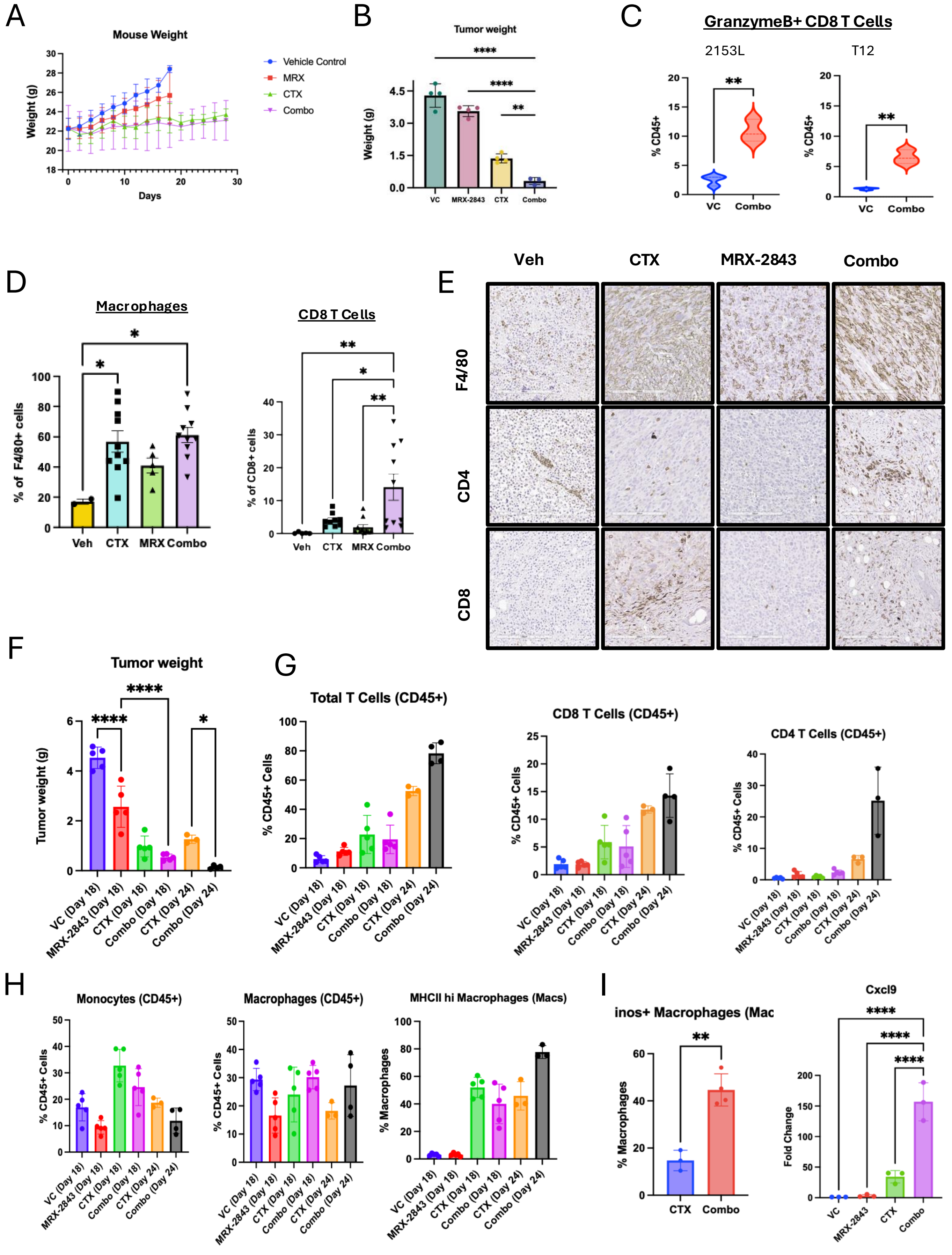

A
Mouse Weight
Weight (g)
Days
Vehicle Control
MRX
CTX
Combo
B
Tumor weight
Weight (g)
VC
MRX-2843
CTX
Combo
C
GranzymeB+ CD8 T Cells
2153L
T12
% CD45+
VC
Combo
D
Macrophages
% of F4/80+ cells
Veh
CTX
MRX
Combo
CD8 T Cells
% of CD8+ cells
E
Veh
CTX
MRX-2843
Combo
F4/80
CD4
CD8
F
Tumor weight
Tumor weight (g)
VC (Day 18)
MRX-2843 (Day 18)
CTX (Day 18)
Combo (Day 18)
CTX (Day 24)
Combo (Day 24)
G
Total T Cells (CD45+)
% CD45+ Cells
CD8 T Cells (CD45+)
CD4 T Cells (CD45+)
H
Monocytes (CD45+)
Macrophages (CD45+)
MHCII hi Macrophages (Macs)
% Macrophages
I
inos+ Macrophages (Mac
Cxcl9
Fold Change

**Supplemental Figure 3. Temporal TME cell populations predict response. CD4 infiltration predicts LTR.** (A) Mouse weight of 2153L tumor bearing mice across treatment groups. (B) Tumor weight at day 18 across treatment groups. (C) Vln plots showing increased GranzymeB$^{+}$CD8$^{+}$ T Cells in combination treated 2153L tumors. (D) Quantification of IHC analyses. (E) Representative IHC images. (F) Bar plots comparing tumor weight at day 18 T12 and day 24 T12 tumors. (G) Bar plots of flow cytometry analyses comparing day 18 T12 TME to day 24 T12 TME. (H) Bar plot of protein expression of iNOS and RNA expression of CXCL9 in day 24 combination treated T12 tumors.

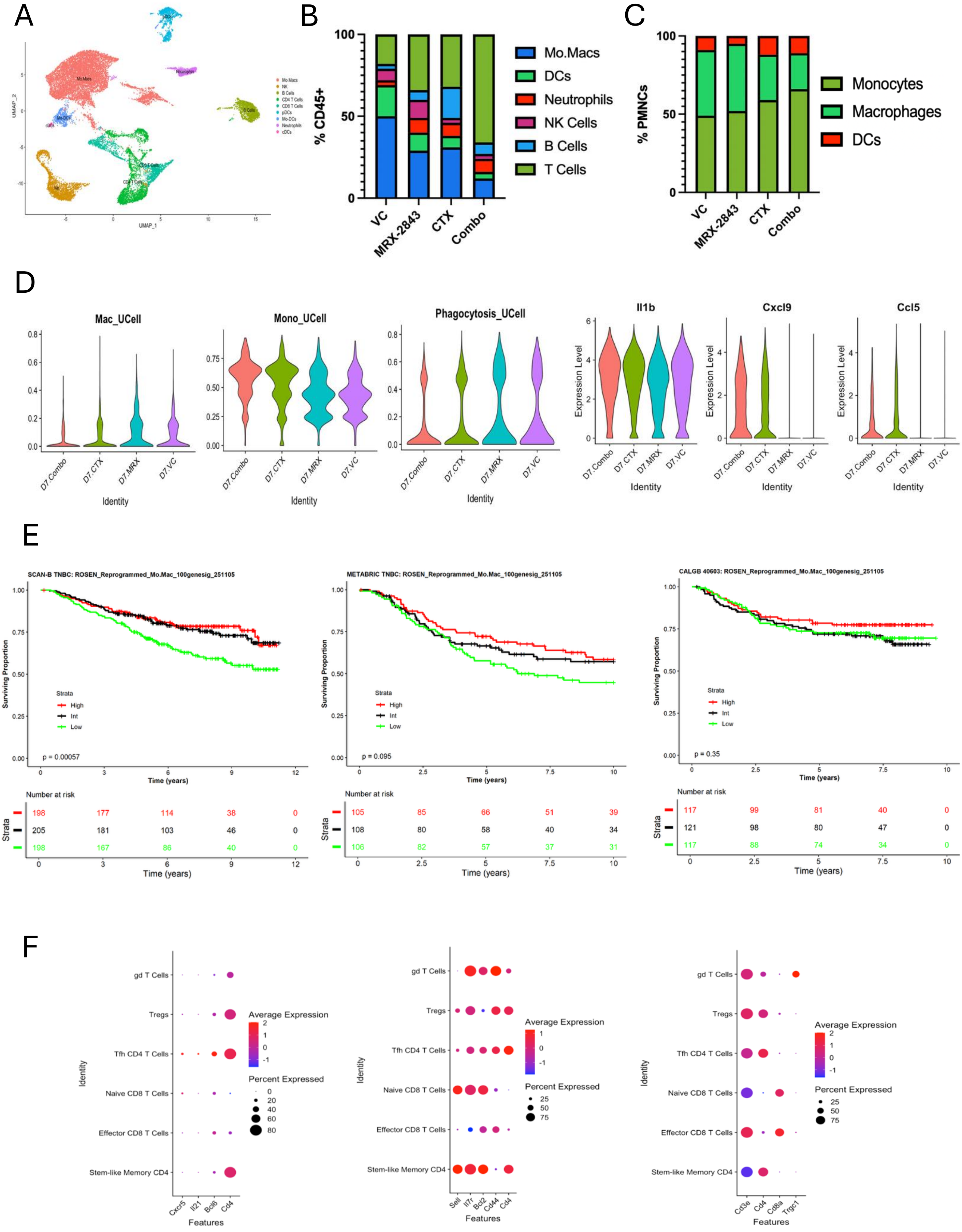
A
UMAP_1
UMAP_2
Mo.Macs
NK
B Cells
CD4 T Cells
CD8 T Cells
pDCs
Mo-DCs
Neutrophils
cDCs
B
% CD45+
VC
MRX-2843
CTX
Combo
Mo.Macs
DCs
Neutrophils
NK Cells
B Cells
T Cells
C
% PMNCs
VC
MRX-2843
CTX
Combo
Monocytes
Macrophages
DCs
D
Mac_UCell
Mono_UCell
Phagocytosis_UCell
Il1b
Cxcl9
Ccl5
Expression Level
D7.Combo
D7.CTX
D7.MRX
D7.VC
Identity
E
SCAN-B TNBC: ROSEN_Reprogrammed_Mo.Mac_100genesig_251105
METABRIC TNBC: ROSEN_Reprogrammed_Mo.Mac_100genesig_251105
CALGB 40603: ROSEN_Reprogrammed_Mo.Mac_100genesig_251105
Surviving Proportion
Strata
High
Int
Low
p = 0.00057
p = 0.095
p = 0.35
Time (years)
Number at risk
198 177 114 38 0
205 181 103 46 0
198 167 86 40 0
105 85 66 51 39
108 80 58 40 34
106 82 57 37 31
117 99 81 40 0
121 98 80 47 0
117 88 74 34 0
F
gd T Cells
Tregs
Tfh CD4 T Cells
Naive CD8 T Cells
Effector CD8 T Cells
Stem-like Memory CD4
Identity
Features
Average Expression
Percent Expressed
Cxcr5
Il21
Bcl6
Cd4
Sell
Il7r
Bcl2
Cd44
Cd3e
Cd8a
Trgc1

**Supplemental Figure 4. Single Cell analysis of TME reveals Mo.Mac signature and SLM CD4 T Cells coordinate response.** (A) UMAP of all CD45$^{+}$ cells across day 7 treatment datasets. (B) Bar plot of cell percentages in dataset. (C) Bar plot of monocyte (LY6C+F4/80$^{-}$), macrophage(F4/80$^{+}$), DC (SiglecH$^{+}$ or CD209a$^{+}$) cell percentages of PMNCs. (D) UCell signature vln plots across treatments. (E) KM plots of CXCL9$^{+}$ Mo.Mac signature in SCAN-B, METABRIC, and CALGB TNBC datasets. (F) Dot plots of T Cell marker genes.

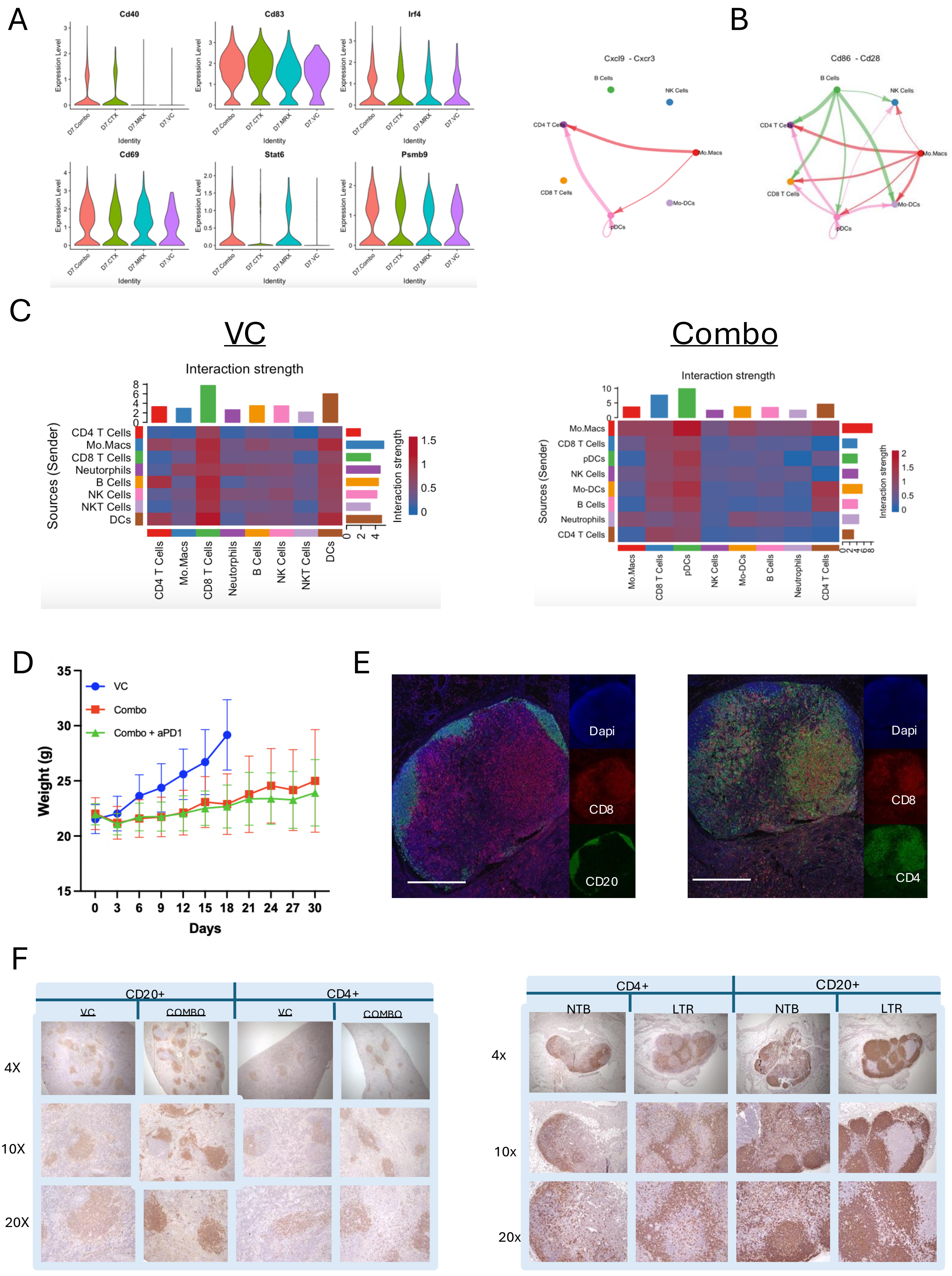
A
Cd40
Cd83
Irf4
Cd69
Stat6
Psmb9
Expression Level
D7.Combo
D7.CTX
D7.MRX
D7.VC
Identity
B
Cxcl9 - Cxcr3
Cd86 - Cd28
B Cells
NK Cells
CD4 T Cells
Mo.Macs
CD8 T Cells
Mo-DCs
pDCs
C
VC
Combo
Interaction strength
Sources (Sender)
CD4 T Cells
Mo.Macs
CD8 T Cells
Neutrophils
B Cells
NK Cells
NKT Cells
DCs
pDCs
Mo-DCs
D
VC
Combo
Combo + aPD1
Weight (g)
Days
E
Dapi
CD8
CD20
CD4
F
CD20+
CD4+
VC
COMBO
NTB
LTR
4X
10X
20X
4x
10x
20x

**Supplemental Figure 5. CD4 lymphocytes and B Cells have increased signaling and prevalence in combination treated mice.** (A) Vln plot of B cell activation genes. (B) Circus plot of CXCL9-CXCR3 cellchat in combination treated CD45+ cells. (C) Cell chat analysis of all signaling between CD45+ cells. (D) Mouse weight of mice treated with combination or combination + aPD1. (E) Representative IF images of TDLNs in T12 tumor bearing combination treated mice and 2153L tumor bearing combination treated mice at day 30. (F) IHC of spleens at day 18 and TDLN at day 30 of 2153L tumor bearing mice.
